# Chaetognaths exhibit the most extensive repertoire of Hox genes among protostomes

**DOI:** 10.1101/2025.01.31.635887

**Authors:** June F. Ordoñez, Tim Wollesen

**Affiliations:** Unit for Integrative Zoology, Department of Evolutionary Biology, University of Vienna, 1030 Vienna, Austria

## Abstract

The evolutionary origins and body plan diversification within bilaterians hinge on our understanding of conserved developmental gene networks across diverse taxa. While significant advances have been made in elucidating the anterior-posterior (AP) axis patterning across well-studied bilaterian lineages, our understanding of the conservation of AP-patterning gene expression in underexplored protostomes with a phylogenetically informative position remains limited. Chaetognaths, a group of marine invertebrates, form a clade with Gnathifera as a sister to the remaining Lophotrochozoa, occupying an early-diverging branch of the spiralian lineage. Their phylogenetic position provides potentially valuable evolutionary insights into whether the AP patterning reflects conserved bilaterian mechanisms or reveals distinct lineage-specific adaptations. Here, we investigate the expression patterns of anterior nervous system markers (*otx*, *nk2.1*, *six3/6*) and Hox genes in post-embryonic stages of the chaetognath *Spadella cephaloptera* using fluorescence whole-mount *in situ* hybridization. We identify expression domains of anterior-patterning genes in the cerebral ganglion and head structures, consistent with their conserved role in anterior central nervous system (CNS) specification in bilaterians. Additionally, we describe a staggered expression pattern of Hox genes, including previously undescribed central (*Sce-med6*) and posterior class (*Sce-postC* and *Sce-postD*), along the ventral nerve cord (VNC) and post-anal tail. Our results demonstrate that chaetognaths exhibit the most extensive repertoire of Hox genes among protostomes, within metazoans only surpassed by chordates. All AP patterning genes are expressed in a staggered manner, with Hox gene expression absent in the head region. This pattern resembles the conserved expression profile inferred for the last common bilaterian ancestor and is only rudimentarily visible in other spiralians, including annelids and mollusks. Posterior Hox genes including the newly discovered *postC* and *postD* genes are absent in the hitherto investigated gnathiferan sister groups such as rotifers. The absence of a postanal tail in rotifers and other gnathiferans, combined with the expression of posterior Hox genes in the elongated postanal tail region, suggests their involvement in the formation of this unique chaetognath structure. Posterior flexibility of Hox genes, as previously hypothesized for chordates, likely contributed to the formation of the chaetognath tail during the early Cambrian period.

## 2. Introduction

The anterior-posterior (AP) axis is a fundamental feature of bilaterian body plans, established by a suite of highly conserved transcription factors that orchestrate regional identity and developmental programs. Among these, anterior-specific transcription genes, such as *fezf*, *gbx*, *irx*, *nk2.1*, *otx*, *pax2/5/8*, and *six3*, are essential for patterning the anterior body region, while Hox genes play a critical role in defining positional information along the AP axis <u>(e.g. Arenas-Mena et al., 2000; Eriksson et al., 2013; Feuda and Peter, 2022; Fritsch et al., 2015; Hiebert and Maslakova, 2015; Holland et al., 2013; Janssen et al., 2014; Lowe et al., 2003; Marlow et al., 2014; Omori et al., 2011; Redl et al., 2018; Steinmetz et al., 2011)</u>. In addition, anterior-specific genes and Hox genes show conserved AP expression in the central nervous system (CNS) of diverse bilaterians, underscoring their critical role in neural regionalization (Aronowicz and Lowe, 2006; Faltine-Gonzalez et al., 2023; Hejnol and Lowe, 2015; Holland et al., 2013; Jarvis et al., 2012; Philippidou and Dasen, 2013). Despite extensive research in model organisms such as vertebrates, arthropods, and annelids, our understanding of how these genes contribute to AP axis formation across the broader diversity of bilaterians remains incomplete. Comparative studies in underexplored taxa, particularly within Spiralia, are essential to uncover the conserved and lineage-specific mechanisms driving AP axis formation and the evolution of bilaterian body plans (e.g. Martín-Durán et al. 2018).

Chaetognaths (arrow worms), a group of enigmatic marine invertebrates, have been assigned to various phylogenetic positions due to their mosaic of morphological and molecular traits, which have been interpreted as shared with either deuterostomes, protostomes, or neither (Dunn et al., 2008; Giribet et al., 2000; Harzsch and Müller, 2007; Helfenbein et al., 2004; Laumer et al., 2019; Littlewood et al., 1998; Mallatt and Winchell, 2002; Marlétaz et al., 2019, 2006; Matus et al., 2006; Papillon et al., 2004, 2003; Paps et al., 2009; Peterson and Eernisse, 2001; Philippe et al., 2011; Telford and Holland, 1993). In this context, arrow worms may represent a crucial taxon to infer spiralian evolution. Fossil evidence of complete chaetognaths, along with their grasping spines, suggest that they were already occupying ecological niches as bilaterian top-level predators, dating back to the earliest Cambrian (Terreneuvian; Park et al., 2024). This contrasts with other gnathiferan groups, for which only reliable younger fossils have been identified so far. Remarkably, the chaetognath body plan has remained largely unchanged and persists to the present day. Chaetognaths share several developmental and morphological features with rotifers, including the lack of spiral cleavage, the specification of primordial germ cells (PGCs) through preformation, a head with a ciliated organ (corona) and hard, chitinous structures for feeding, a trunk without external motile cilia, a post-anal structure (tail/foot), and the central-post class Hox gene chimera, *medpost* (Fröbius and Funch, 2017). While chaetognaths are suggested to have a close evolutionary relationship to gnathiferans (Fröbius and Funch, 2017), their exact position remains debated, with hypothesis positioning them either within Gnathifera (comprising rotifers, acanthocephalans, gnathostomulids, and micrognathozoans; Marlétaz et al., 2019) or as a sister lineage to the group (Laumer et al., 2019; Park et al., 2024), forming a clade called Chaetognathifera (Bekkouche and Gąsiorowski, 2022; Park et al., 2024). These molecular phylogenomic analyses suggest that chaetognaths and gnathiferans form a group that is a sister lineage to all Lophotrochozoa (Laumer et al., 2019; Marlétaz et al., 2019), thus offering a unique perspective on ancestral spiralian developmental features.

A comprehensive understanding of anterior-posterior (AP) patterning in chaetognaths remains lacking, especially in comparison to other spiralians. Previous studies have examined the Hox complement of *Spadella cephaloptera* and *Sagitta enflata* (Matus et al., 2007; Papillon et al., 2003), revealing a repertoire, although incomplete, closely resembles those of other bilaterians. Papillon and colleagues <u>(2005)</u> have provided the first insight into Hox gene expression, showing that a Hox gene related to paralogous group (PG) 5 or *sex comb reduced* (*scr*) genes exhibits restricted expression in the ventral nerve center (VNC). In contrast, the rotifer *Brachionus manjavacas* possesses a reduced Hox gene complement, notably lacking the posterior-class Hox genes (Fröbius and Funch, 2017). Furthermore, later on their expression has been interpreted as partially staggered (Huan et al., 2020) and limited to the developing nervous system (Fröbius and Funch, 2017), highlighting their role in neural patterning rather than whole-body regionalization. Moreover, anterior-patterning genes *nk2.1* and *pax6* are expressed in the brain of the juvenile rotifer *Epiphanes senta* (Martín-Durán et al., 2018). These findings raise questions about how chaetognath AP patterning genes contribute to AP axis development, particularly considering their phylogenetic position relative to Gnathifera. Unlike rotifers, chaetognaths retain Hox genes from anterior, central, and posterior classes, suggesting that their expression may reflect a conserved bilaterian mechanism. Consequently, expanding investigations into the expression dynamics of both Hox and anterior-patterning genes in chaetognaths is essential for testing this hypothesis. Such studies will also help identify conserved features and lineage-specific adaptations in chaetognaths, further providing deeper insights into their evolutionary and developmental relationships within Spiralia.

Here, we examine the expression profiles of AP patterning genes in the chaetognath *Spadella cephaloptera* during early post-embryonic development, focusing on the expression of *otx*, *nk2.1*, *six3/6*, and the most comprehensive set of Hox genes published for gnathiferans so far. Our findings reveal expression of anterior patterning genes in the head structures, including the developing cerebral ganglion, consistent with their conserved role in the specification of brain and anterior tissues observed across Bilateria. Additionally, we provide an updated Hox gene complement for *S. cephaloptera*, including previously undescribed members, and demonstrate their staggered expression along the AP axis of the VNC and the post-anal tail. These spatially organized expression patterns suggest that chaetognaths retain core bilaterian mechanisms of CNS patterning that have been secondarily lost in numerous other bilaterians studied so far, despite their distinct developmental features and evolutionary position.

## 3. Methods

### 3.1. Identification of gene homologs

AP patterning (*otx*, *six1/3*, *nk2.1*, Hox) genes were retrieved from the *S. cephaloptera* draft transcriptome (Wollesen et al., 2023) using blastx searches (Altschul et al., 1990) against protein sequences from NCBI Genbank non-redundant protein database. Phylogenetic reconstruction was used to determine gene orthology. Reference amino acid sequences from other bilaterians were downloaded from NCBI Genbank (Table S1). Multiple sequence alignments were performed with MAFFT (v7.490) (Katoh, 2005, 2002) implemented in Geneious Prime (v2023.0.1). The resulting multi-sequence alignments were then trimmed with ClipKit (v2.2.3) (Steenwyk et al., 2020). The maximum likelihood (ML) was performed in IQTREE (2.3.6), incorporating ModelFinder for amino-acid substitution model selection and conducting 1,000 ultrafast bootstrap replicates (Hoang et al., 2018; Kalyaanamoorthy et al., 2017; Minh et al., 2020) and alongside SH-aLRT test replicates for Hox genes (Guindon et al., 2010). All parameters used in phylogenetic analyses are indicated in Supplementary File 1 Table S1. Newly obtained *S. cephaloptera* sequences were submitted to GenBank (Accession number: XXXX).

### 3.2. Animal collection

Live adult specimens of *Spadella cephaloptera* (Busch, 1851) were collected in June 2024 during low tide from the intertidal zone near Roscoff, France (48°43’47.2"N 3°59’12.2"W) and were housed in aquaria in the aquatic research facility of the University of Vienna. The aquaria contain a combination of natural and artificial seawater, with a temperature and salinity of 14°C and 35**‰**, respectively. Approximately 30 sexually matured individuals were collected from the aquaria and transferred to plastic Petri dishes (14.1 cm radius), which were refreshed with natural sea water every day. The animals were fed with *Artemia sp.* nauplii until they laid eggs. Embryos and hatchlings (8 - 20 hours post-hatching) were manually collected. Additionally, some hatchlings were allowed to develop to early juveniles (7-10 days post-hatching) before they were sampled.

All specimens were fixed in 4% paraformaldehyde in a buffer containing 0.1 M MOPS pH 7.4, 2 mM EGTA, 1 mM MgSO_4,_ 2.5M NaCl. The specimens used for whole-mount RNA Fluoresce In situ Hybridization (RNA-FISH) were fixed overnight (14 - 18 hours) while for whole-mount Hybridization Chain Reaction (HCR) – RNA-FISH (Choi et al., 2018; Tsuneoka and Funato, 2020; Wang et al., 2024), they were fixed for only 1 – 1.5 hours. Following fixation, the specimens were washed three times for five minutes each in PBTw (1x PBS, pH 7.5 with 0.1% Tween-20), washed three times in ice-cold methanol for 10 minutes each, and then stored in 100% methanol at −20°C until use.

### 3.3. Sequencing, riboprobe synthesis, and RNA fluorescent *in situ* hybridization

RNA extracted from hatchlings and adults using RNAqueous™-Micro Total RNA Isolation Kit (Invitrogen GmbH, Karlsruhe, Germany) was used to generate a cDNA library with First Strand cDNA Synthesis Kit for RT-PCR (AMV) (Roche Molecular Biochemicals, Vienna, Austria). Riboprobes were generated either directly from gene fragments amplified by PCR or from plasmid cloning. Gene-specific primer pairs for PCR-derived riboprobes were designed with a T7 sequence of the 5’ end of the reverse primer and resulting products were purified with QIAquick PCR Purification Kit (Qiagen Vertriebs GmbH, Vienna, Austria). For cloning, gene fragments amplified with PCR were cloned into pGEM-T Easy vectors (Promega, Mannheim, Germany), transformed into competent *Escherichia coli* cells, and then purified with QIAprep® Miniprep (Qiagen Vertriebs GmbH, Vienna, Austria). The purified fragments from both methods were sent to Microsynth Austria for Sanger sequencing to validate their specificity. Primer sequences and PCR conditions are listed in Supplementary File 1 Table S2. Digoxigenin (DIG)-labeled sense riboprobes from the PCR product and linearized DNA were generated using the T7 polymerase and digoxigenin RNA labeling mix kit (Roche Diagnostics, Mannheim, Germany). Total riboprobe concentrations were estimated using NanoDrop 2000 spectrophotometer (ThermoFisher Scientific, Waltham, MA). The traditional whole-amount fluorescent *in situ* hybridization (referred to as RNA-FISH in the succeeding sections) experiments on hatchlings followed the procedure previously described in (Ordoñez and Wollesen, 2024; Wollesen et al., 2023). HCR RNA-FISH (Choi et al., 2018, 2014) was carried out with genes (*Sce-hox2*, *Sce-medpost*, *Sce-postB*, *Sce-postC*, *Sce-postD*, *Sce-nk2.1,* and *Sce-otx*) for which riboprobes failed to work with the conventional RNA-FISH. Probes for each gene were designed using the program *insitu_probe_generator.py* (https://github.com/rwnull/insitu_probe_generator; (Kuehn et al., 2022) and were purchased from Integrated DNA Technologies (München, Germany). Hairpins were purchased from Molecular Instruments, Inc. (California, USA). HCR RNA-FISH was performed following (Choi et al., 2016). Information on the probes, hairpins, and their working concentrations are indicated in the Supplementary File 1 Table S3. Specimens were incubated in Vectashield (Vector Laboratories Burlingame, CA) for 20 minutes before mounting with 80% 2,2’-thiodiethanol in PBS. Specimens were scanned with Leica TCS SP5 (Leica Microsystems, Heidelberg, Germany) confocal laser scanning microscope. Image adjustments (brightness and contrast) and Z-stacks of all confocal scans were performed with Fiji (Schindelin et al., 2012). IMARIS (v 3.4.1) was also used to capture images of the orthogonal sections (lateral and transverse planes). Figure assembly and schematic drawings based on post-processed images were made with Inkscape (https://inkscape.org).

## 4. Results

### 4.1. Gene orthology and expression patterns of anterior-patterning genes

Three candidate anterior-patterning genes were identified and characterized to examine their patterns of expression and potential roles during AP patterning of the CNS in *S. cephaloptera*. Single-copy gene orthologs of *otx, nk2.1*, and *six3/6* from the *S. cephaloptera* draft transcriptome were identified with blast search and supported by their discrete grouping with known protein sequences from representative bilaterians as inferred from the ML phylogenetic tree [Supplementary File 1 Table S1 and Supplementary File 2 Fig. S1 and S2; See <u>Ordoñez and Wollesen (2024)</u> for *Sce-nk2.1* orthology assignment].

#### Otx expression

In hatchlings, *Sce-otx* is broadly expressed in the head, marking the cerebral ganglion with the strongest signal concentrated in its anterior-most region (Fig. 1b). In addition, *Sce-otx* expression is detected in the presumptive vestibular ganglia (Fig. 1b’), head epidermis (Fig. 1b’ – b’’’), cephalic adhesive papillae (white arrowhead; Fig. 1b, b’’), and around the future mouth opening (asterisk, Fig. 1b, b’’). In the lateral somata clusters of the VNC, *Sce-otx* is concentrated in the ventral region of the anterior and posterior tips and shows a bilateral streak pattern along the anterior inner cells (Fig. 1b, b’). The primordial germ cells are also positive for *Sce-otx* (Fig. 1b’’’). In the early juvenile, *Sce-otx* is also widely expressed throughout head (Fig. S7b – b’’’). *Sce-otx* transcripts are observed in the brain as concentrated patches in the anterior tip and lateral regions of the anterior and posterior brain (dotted outline; Fig. S7b’), corona ciliata (dashed outline; Fig. S7b’, b’’’), eyes (circles; Fig. S7b’), perioral epidermis (red arrowheads; Fig. S7b’’, b’’’), hood (Fig. S7b’’’), and cephalic epidermis (Fig. S7b, b’’’). In the trunk, *Sce-otx* is expressed in the lateral somata clusters, specifically in the anterior tips (Fig. S7b, b’’). *Sce-otx* transcripts are also present in the proliferating germ cells (dotted outline, Fig. S7b’’’’).

**Figure 1.**
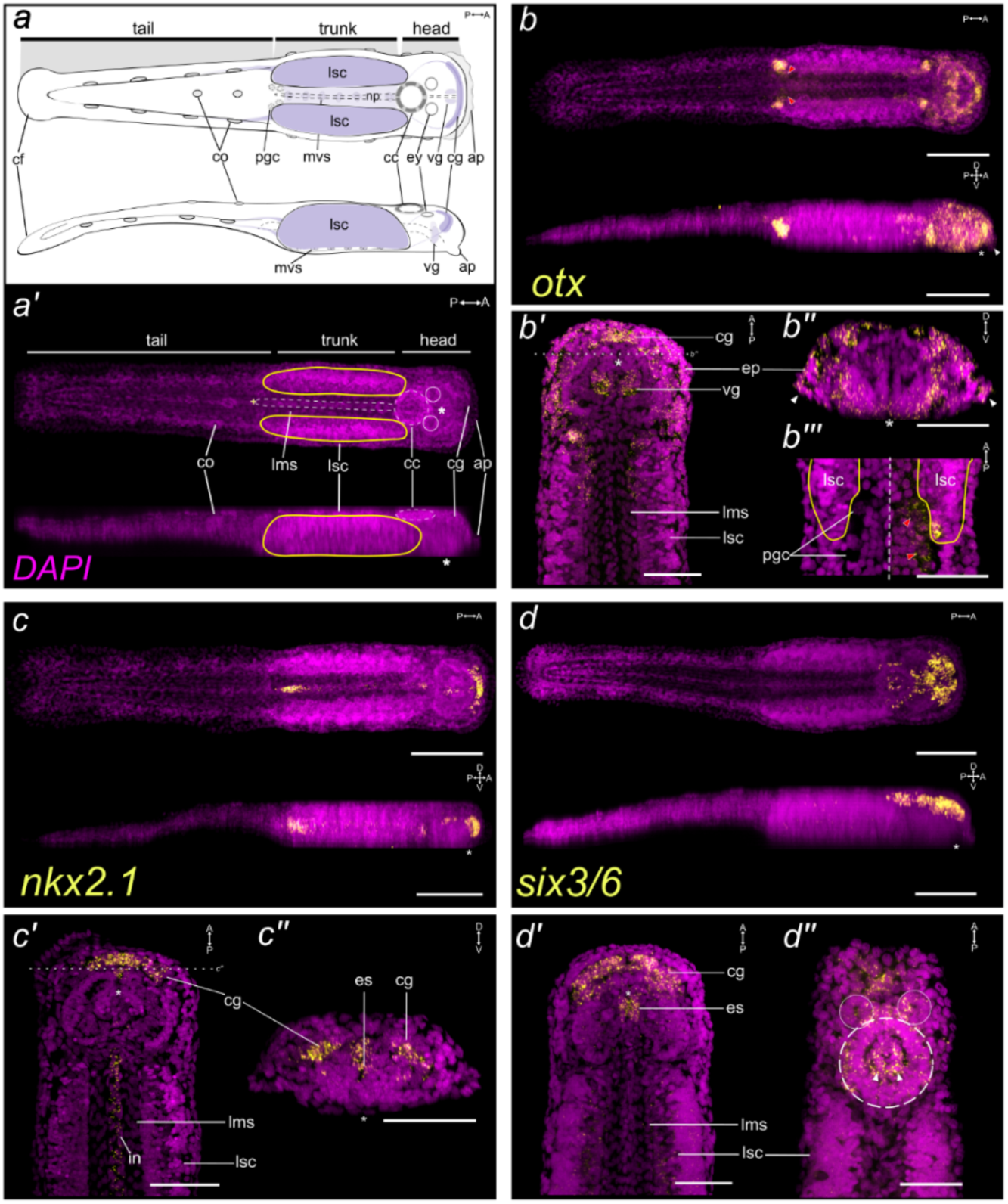
Expression pattern of *Sce-otx, Sce-nk2.*1, and *Sce-six3.6* in *S. cephaloptera* hatchling (1 dph). Gene transcripts (yellow) are visualized with AlexaFluor-647 (b, c) or AP-Fast Blue (d) and cell nuclei with DAPI (purple). General morphology of a *S. cephaloptera* hatchling in schematic drawing (a) and maximum intensity projection of the whole animal in DAPI (a’’). Top and bottom figures are coronal and lateral view, respectively. (b – b’’’) *Sce-otx* expression pattern. (b’) Coronal profile exposing the presumptive vestibular ganglion (*vg*). (b’’) Transverse profile behind the anterior portion of the cerebral ganglion (*cg*). Posterior parts of the cephalic adhesive papillae (*ap*) are indicated in white arrowheads. (b’’’) Higher magnification of the primordial germ cells (*pgc*; red arrowheads) in DAPI channel (left side) and both channels (right). (c – c’’) *Sce-nk2.1* expression pattern. (c’) Coronal profile along the gut. (c’’) Transverse profile behind the anterior portion of the cerebral ganglion. (d – d’’) *Sce-six3/6* expression pattern. (d’, d’’) Coronal profile along the intestine (*in*, d’) and the dorsal-most portion showing the eyes (solid circles) and corona ciliata (dashed circle, d’’). Scale bars: 100 μm, except panels *b’’*, *b’’’* and *c’* (50 μm). Asterisk indicates the position of the mouth opening. Orientation of specimens is in the top right corner of each panel. ap, cephalic adhesive papillae; cc, corona cilata; cf, caudal fin; cg, cerebral ganglion; co, ciliary tuft/fence organ; es, esophagus; in, intestine; lms: longitudinal mucles somata; lsc, lateral somata clusters; mvs, medioventral somata clusters; pgc, primordial germ cells; vg, vestibular ganglion.

#### Nk2.1 expression

At hatching, *Sce-nk2.1* expression appears as a semi-arch, marking the anterior part of the developing brain (Fig. 1c, c’’). It also outlines the digestive track, from the esophagus to the posterior gut, which shows the strongest signal intensity (Fig. 1c). The expression domains of *Sce-nk2.1* in the early juvenile is observed in the anterior tip of the anterior brain and in patches of neurons in its posterior domain (Fig. S7c, c’’). *Sce-nk2.1* remains active in the cells of the digestive tract at this stage (Fig. S7c, c’’, c’’’).

#### Six3/6 expression

*Sce-six3/6* is mostly expressed in the head of hatchlings (Fig. 1d). It is observed in the cerebral ganglion, eyes, corona ciliata, and dorsoanterior epidermal cells of the head (Fig. 1d – d’’). The cells of the esophagus are also positive for *Sce-six3/6* (Fig. 1b’). A faint expression of the transcript can be observed in the lateral somata clusters. *Sce-six3/6* in the early juveniles show most prominent expression around the longitudinal midline of the brain (dotted outline, Fig. S7d, d’). A less intense expression is also observed in the corona ciliata (dashed outline, Fig. S7d’), eyes (circles, Fig. S7d’), vestibular ganglia, esophageal ganglia (Fig. S7d’’), and the lateral somata clusters (Fig. S7d’’’).

### 4.2. *Spadella* Hox gene complement and orthology

Chaetognaths are currently known to have ten Hox genes representing five paralogous groups: anterior class Hox genes, *hox*1 (PG 1-2), and *hox*3 (PG 3); central class *hox4* and *hox5* (PG 4-5) and *hox6*, *hox7*, and *hox8 (*PG 6-8); *medpost*; and posterior class *postA* and *postB* (PG 9-14) (Matus et al., 2007; Papillon et al., 2003). We surveyed the *S. cephaloptera* draft transcriptome and identified orthologs of the previously reported Hox genes in chaetognaths using blast searches and ML-based phylogenetic approaches (Fig. S1). Four previously undescribed Hox genes in chaetognaths are identified: *Sce-hox2*, *Sce-med6*, *Sce-postC*, and *Sce-postD*. Orthology analysis assigns *Sce-hox2* to *hox*2/*proboscipedia* (*pb*) cluster with SH-aLRT support (SS) and ultrafast bootstrap support (UFBS) of 79.9% and 82%, respectively (Fig. S3), and it contains five amino-acid residues unique to PG2: R (position 2), L (4), N (10 and 23), and V (45) (Fig. S4). Based on the homeodomain and diagnostic amino-acid motifs, *Sce-med6* displays high similarity with central Hox genes, sharing all eleven diagnostic amino acids (Papillon et al., 2003) in its homeodomain: Q (6), T (7), R (10), LTR(R/K)RRI (26–32), and E (59) (Fig. S5). Notably, *Sce-med6* clustered with *hox8*/*lox2*/l*ox4*/*ubx*/*abdA* sequences (SS: 80.4% and UFBS: 92%; Fig. S3). While orthology analysis shows *Sce-med6* close relationship with to chaetognath *medpost* (SS: 73.7% and UFBS: 99%), all diagnostic posterior class amino acid residues are absent except for the residue Y at position 20 (Fig. S5).

A blastp search of *Sce-postC* and *Sce-postD* coding sequence against the nr database reveals general similarity to the vertebrate *hox9/10* and arthropod abdominal (*abdB*), respectively. Orthology analysis cluster them to the *post1/hox11 – 13* group of the posterior Hox class (PG 9-13; SS: 83.8% and UFBS: 54%) (Fig. S3). Examining their amino acid sequence shows that the diagnostic posterior class residues (Matus et al., 2007; Papillon et al., 2003), K3 and R18 are present in both genes, while V (21) is only found in *Sce-postD* (Fig. S6). Also, *Sce-postC* shares another posterior class residue Y (20) together with *Sce-postA* and *Sce-postB* (Fig. S6), which is considered as a spiralian *post1* signature (Matus et al., 2007).

### 4.3. Expression patterns of *Spadella* Hox genes

Fluorescent in situ hybridization (FISH) analysis revealed a staggered spatial expression pattern of Hox genes along the anterior–posterior (AP) axis in early post-embryonic stages of *S. cephaloptera* (Fig. 2). In the hatchling, *Sce-Hox1* is expressed two bilateral streaks in the inner region of the anterior lateral somata clusters (Fig. 2b) and in the longitudinal muscle of the trunk, while *Sce-hox2* spans most of the lateral somata clusters except in the anterior and posterior tips (Fig. 2c). Similarly, *Sce-hox3* is expressed as two distinct vertical bands in the VNC (Fig. 2d), overlapping with *Sce-hox2* in the mid-region but extending only slightly before the anterior and posterior boundaries of the *Sce-hox2* domain. Both genes overlap with the expression domains of central class Hox genes. *Sce-hox4*, *Sce-hox5*, *Sce-hox6*, and *Sce-hox7* form horizontal bands in the medial lateral somata clusters, with trailing anterior (*Sce-hox7*) and posterior (*Sce-hox4*, *Sce-hox5*, *Sce-hox6*, and *Sce-hox7*) expression signals, indicating overlapping transcriptional domains (Fig. 2e – h). *Sce-hox8* is expressed in a broader domain in the posterior VNC, predominantly spanning the posterior half (Fig. 2i). *Sce-hox5, Sce-hox6*, *Sce-hox7*, and *Sce-hox8* are also observed in the medioventral somata clusters within the middle position of the VNC (red arrowheads, Fig. 2f – i, f’’ – i’’). *Sce-hox8* is also expressed in the longitudinal muscle of the posterior trunk (white arrowheads, Fig. 3i, i’’). *Sce-med6* also shows as a horizontal band within the posterior half of the lateral somata clusters while *Sce-medpost*, displaying a triangular-shaped pattern, occupies a larger region in its posterior end (Fig. 2j – k). The posterior class Hox genes, *Sce-postA*, *Sce-postC, and Sce-postD* are restricted to the posterior tip, with the latter two extending further than *Sce-postA* (Fig. 2l – n). *Sce-hox8*, *Sce-med6*, *Sce-postC, and Sce-postD* are also expressed in the hindgut. Notably, *Sce-postB* shows no detectable expression across all developmental stages examined (data not shown). In the early juvenile, almost similar expression pattern in the VNC is also observed (Fig. S8)

**Figure 2.**
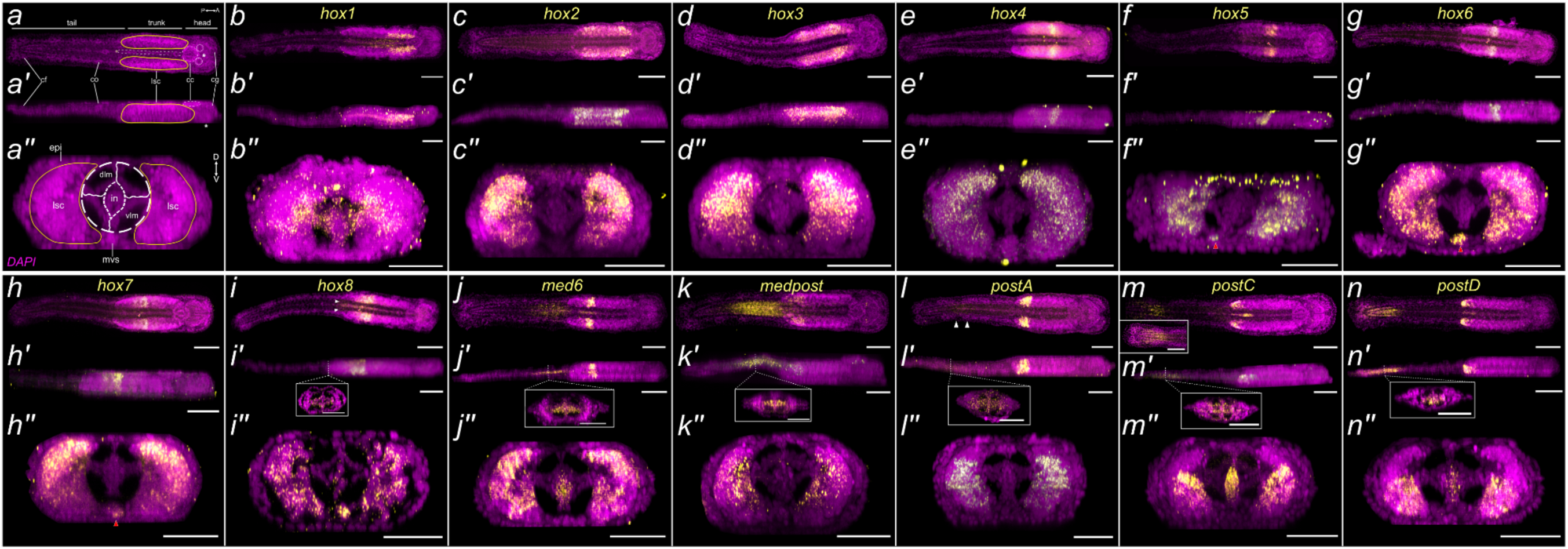
Expression pattern of Hox genes in *S. cephaloptera* hatchling (1 dph). Gene transcripts (yellow) are visualized with AlexaFluor-647 (c, j, k, m, n) or AP-Fast Blue (b, d – i, l) and cell nuclei with DAPI (purple). (a) General morphology of a *S. cephaloptera* hatchling in maximum intensity projection of the whole animal in DAPI, with panel *a* and *b* showing coronal view and lateral view, respectively. The eyes are encircled and gut in dashed structure in between the lateral somata clusters (lsc). The position of the mouth and anal opening are indicated in asterisk and cross, respectively. (a’’) shows the transverse section of the trunk. Inset in *i’ – n’* is a transverse profile along the tail. Red arrowheads (f – i) indicate medioventral somata cluster that expresses Hox gene while white arrowhead (i, l) indicate expression domain. Inset in *m* shows a clearer DAPI view of the caudal region of the tail. Scale bars: 100 μm, except panels *b’’ – n’’* (50 μm) and all insets (25 μm). Orientation of specimens is in the top right corner of each panel. cc, corona cilata; cf, caudal fin; cg, cerebral ganglion; co, ciliary tuft/fence organ; dlm, dorsal longitudinal muscle; in, intestine; lsc, lateral somata clusters; mvs, medioventral somata clusters; vlm, ventral longitudinal muscle.

**Figure 3.**
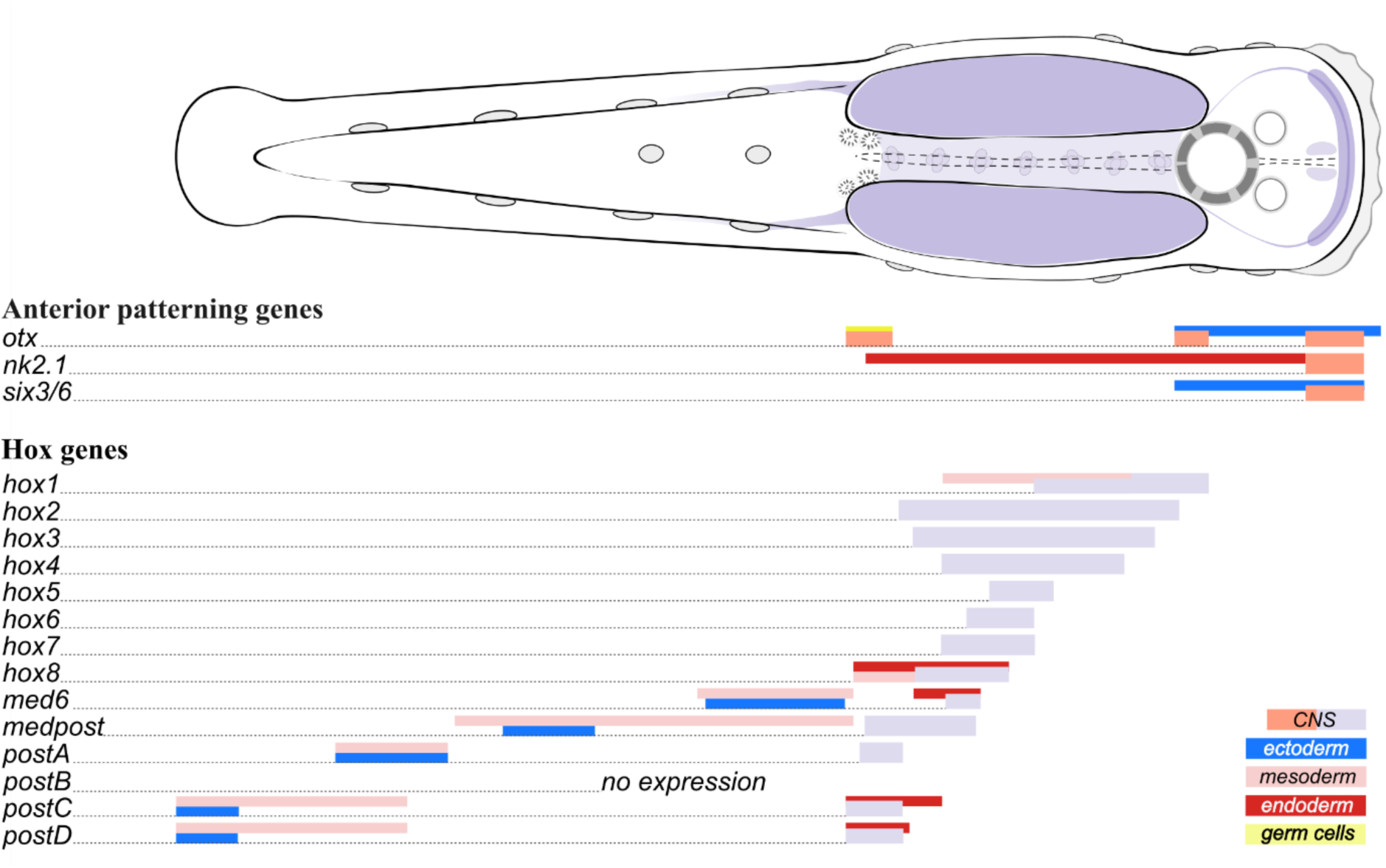
Schematic representation of the spatial expression patterns of anterior-posterior (AP) patterning genes in *S. cephaloptera* hatchling (1 dph). The anterior patterning genes are expressed in the head region, including the cerebral ganglion and other anterior structures, while Hox genes exhibit a staggered expression pattern along the ventral nerve cord (VNC) and post-anal tail. Bars represent approximate expression domains with different bar colors denoting the germ-layer origin of the tissues.

In hatchlings and early juveniles, posteriorly expressed Hox genes in the VNC exhibit localized and sequential patterns in the tail (Fig. 3i – n; Fig. S8j, k, m, n). *Sce-med6* shows in the anterior tail (Fig. 3j, j’; Fig. S8j), *Sce-medpost* covers the whole anterior half (Fig. 3k, k’; Fig. S8k), *Sce-postA* occupy the pre-posterior end (Fig. l, l’), and *Sce-postC* and *Sce-postD* are restricted to posterior tail, including the caudal fin (Fig. 3m, n, m’, n’; Fig. S8m, n). No expression of *Sce-postA* was detected in the tail of early juvenile. Hox expression in the tail is observed mainly in the cells of the longitudinal muscles and slightly in the epidermal, fin, mesoderm-derived (i.e. lateral, mesenterial, and medial) cells (Fig. 3i – m; Fig. S8j, k, m, n).

## 5. Discussion

Our study explores the molecular basis of anterior-posterior axis formation in chaetognaths, a group occupying a distinctive phylogenetic position within Spiralia. Using *Spadella cephaloptera* as a model, we characterized the expression profiles of key conserved transcription factors involved in AP patterning, including *otx*, *nk2.1*, *six3/6*, Hox genes, and reveal their potential roles during post-embryonic development. These findings provide critical insights into how chaetognath developmental programs relate to those of other spiralians and contribute to our understanding of conserved and lineage-specific mechanisms of AP patterning in Bilateria.

### 5.1. Expression pattern of head-specific genes

The expression patterns of *otx*, *six3/6*, and *nk2.1* in *S. cephaloptera* exhibit similar domains in the anterior regions of other bilaterians (Fig. 3; Andrikou et al., 2019; Eriksson et al., 2013; Martín-Durán et al., 2015; Omori et al., 2011; Redl et al., 2018; Steinmetz et al., 2010; Wollesen et al., 2017).

In hatchlings, *Sce-otx* and *Sce-six3/6* are broadly expressed in the head, aligning with their conserved role in specifying anterior structures across diverse bilaterian taxa (Andrikou et al., 2019; Gonzalez et al., 2017; Hiebert and Maslakova, 2015; Luo et al., 2017; Oliver et al., 1995; Omori et al., 2011; Redl et al., 2018; Santagata et al., 2012; Steinmetz et al., 2010). Interestingly, in many bilaterians, including annelids, molluscs, arthropods, enteropneusts, and vertebrates, *six3/6* expression typically occupies the anterior-most domain, with an *otx-*expression domain positioned posterior to it (Holland et al., 2013; Kaul-Strehlow et al., 2017; Marlow et al., 2014; Redl et al., 2018; Steinmetz et al., 2010; Vöcking et al., 2015). In *Spadella*, however, the expression domains of *Sce-otx* and *Sce-six3/6* overlap at the anterior-most tip of the head. In addition, *Sce-otx* is consistently expressed in the anterior-most region in different embryonic stages (data not shown), indicating that anterior expression domain of *Sce-otx* is conserved between various developmental stages. This observation raises the possibility that the strict anterior-posterior segregation of *six3/6* and *otx* may not be a universal feature of bilaterian development or the patterning could either be modified or lost in chaetognaths.

*Sce-six3/6* expression in the eyes and *Sce-otx* expression in the cells around the oral region is also similar to the observations in many bilaterians (Bruce and Shankland, 1998; Carl et al., 2002; Hinman et al., 2003a; Loosli et al., 1998; Ogura et al., 2013; Oliver et al., 1995; Steinmetz et al., 2010; Vöcking et al., 2015). Also, expression domains in other non-neural anterior structures are observed in the ciliary tuft organs, ciliary fence receptors, corona ciliata, and cephalic adhesive papillae, which are morphological autapomorphies of chaetognaths, reflecting their broader roles in addition to anterior neural patterning and specification. Notably, *Sce-otx* expression is not restricted to the neuroectodermal and ectodermal derivatives, as it also extends to the primordial germ cells. This divergence from the canonical *otx* expression generally observed in bilaterians may reflect functional co-option of *Sce-otx* in the specification of the germ cells of *Spadella*. Although infrequently observed in other spiralians, *otx* is, for example, expressed in some posterior segments and the hindgut of the annelid *Capitella teleta* (Boyle et al., 2014).

The localization of *nk2.1* in the developing cerebral ganglion of *Spadella* hatchlings parallels its patterning in the brain of many bilaterians (Boyle et al., 2014; Harfe and Fire, 1998; Hejnol and Martindale, 2008; Martín-Durán et al., 2018; Takacs et al., 2002; Venkatesh et al., 1999; Zaffran et al., 2000). The cerebral ganglion of the *Spadella* hatchling shows a medial expression of *Sce-nk2.1*, while *Sce-pax6* marks the lateral regions (Ordoñez and Wollesen, 2024). This mediolateral patterning of *nk2.1* and *pax6* in the forebrain has been hypothesized as an ancestral bilaterian feature due to their similar expression in the vertebrate and annelid nervous system (Corbin et al., 2003; Murakami et al., 2001; Tessmar-Raible et al., 2007). While planarians (Currie et al., 2016; Scimone et al., 2014) and the nemertean *Lineus ruber* (Gąsiorowski et al., 2021; Martín-Durán et al., 2018) do not display this expression pattern, other spiralians exhibit the subdivision, including rotifers (Martín-Durán et al., 2018), brachiopods (Martín-Durán et al., 2018; Santagata et al., 2012), and other annelids (Boyle et al., 2014; Klann and Seaver, 2019). Overall, the expression data suggests that *Sce-nk2.1* functions during brain specification and that *Spadella*, even with morphologically simplified brain at hatching, likely retained the ancestral mediolateral spatial division of *nk2.1* and *pax6*.

*Sce-nk2.1*expression in the digestive tract of the hatchling also aligns with its function in gut specification and development across bilaterians (Boyle et al., 2014; Harfe and Fire, 1998; Martín-Durán et al., 2018; Takacs et al., 2002; Venkatesh et al., 1999; Zaffran et al., 2000). *Sce-nk2.1* exhibits a distinct region-specific expression, with higher intensities in the foregut and hindgut, similar to the observations in annelids (Boyle et al., 2014; Kerbl et al., 2016). Other spiralians, including the fruit fly *Drosophila* (Zaffran et al., 2000), the nematode *Caenorhabditis elegans* (Harfe and Fire, 1998), and the gastropod *Haliotis rufescens* (Dunn et al., 2007), only show *nk2.1*/*scarecrow* (*scro*)/*ceh-24* expression in the foregut, while it is expressed in all regions of chordate gut (Ristoratore et al., 1999; Takacs et al., 2002; Venkatesh et al., 1999). This implies that *nk2.1* has a conserved role in defining gut compartments and potentially regulating regional differentiation along the anterior-posterior axis. In addition, *nk2.1*, *foxA* and *otx* are key transcription factors with a conserved role during endomesoderm specification and gut formation in Cnidaria and Bilateria (Hejnol and Martindale, 2008; Omori et al., 2011; Perry et al., 2015). *Sce-foxA* is also expressed in the gut of the hatchling (Ordoñez and Wollesen, 2024), but *Sce-otx* is notably absent in the endomesoderm of the gastrula stages (data not shown) and the gut of hatchling. This suggests that *Spadella* has likely retained only a subset of the core gene regulatory mechanisms associated with endoderm specification and the development of the digestive tract.

### 5.2. New Hox genes expands the repertoire in chaetognaths

Chaetognaths possess a complete set of Hox genes. Our findings reveal 14 Hox genes of *S. cephaloptera* from all paralogous groups, including new members of the central and posterior Hox groups, expanding the known Hox gene complement in chaetognaths. Previous investigations on chaetognath Hox complement only unambiguously identified ten Hox genes: six in *S. cephaloptera* (Papillon et al., 2003) and nine in *Flaccisagitta enflata* (Matus et al., 2007), with both recovering *hox3*, *hox6*, *hox7*, *hox8*, and *medpost* orthologs. While we identified homologs of the ten reported Hox genes, we also recovered unique transcripts of Hox genes not previously described for chaetognaths. Among these is *Sce-hox2*, which belongs to the PG2 group based on ortholog and diagnostic amino acid residue analyses. This finding supports the presence of anterior class Hox genes, including PG1, in chaetognaths, highlighting their conserved complement of anterior Hox genes. Another previously uncharacterized Hox gene in *S. cephaloptera* is *Sce-med6*. Initially, its sequence suggests an orthology to ecdysozoan *ubx*/*abdA* and lophotrochozoan *lox2*/*lox4* genes, as it possesses all the central class diagnostic amino acids (Papillon et al., 2003) and four of the seven residues in the Ubd-A parapeptide motif (xxIxELN; Fig. S5). These features are similar to chaetognath *hox8* except for the configuration of its parapeptide motif (QxIxEM/Lx; <u>Matus et al., 2007 and this study</u>). However, the presence of a posterior class-specific diagnostic residue (tyrosine at homeodomain position 20) suggests *Sce-med6* similarity to *medpost*, a Hox gene possessing both central and posterior class diagnostic residues that is only identified in chaetognath and rotifers thus far (Fröbius and Funch, 2017; Matus et al., 2007). In addition, this assumption is supported by moderate to good branch support values, clustering *Sce-med6* with *medpost* genes of chaetognaths (SS: 73.7% and UFBS: 99%). Moreover, *Sce-med6* is expressed in the tail together with *medpost* and posterior-class Hox genes, further raising the possibility that has indeed a posterior-class Hox signature.

Matus and colleagues (2007) hypothesized that the last common ancestor of protostomes possessed a central-class Hox gene with the UbdA parapeptide, which subsequently underwent independent duplications and divergence occurred in Ecdysozoa and Lophotrochozoa, giving rise to *ubx*/*abdA* and *lox2*/*lox4*, respectively. In this framework, after splitting with lophotrochozoans, lineage-specific modifications within chaetognaths resulted in partial UbdA signal observed in *hox8* (Matus et al., 2007). The identification of *Sce-med6*, another chaetognath Hox that also possesses a divergent UbdA motif, raises the possibility that the inherited ancestral UbdA-containing Hox may have also duplicated and undergone secondary changes in chaetognaths.

Additionally, we identified two more Hox genes that exhibit posterior class Hox genes features. Interestingly, *Sce-postC* and *Sce-postD* group with the *post1*/*hox13* cluster (SS: 83.8% and UFBS: 54%) while *Sce-postA* and *Sce-postB* with *post2*/*abdB* cluster (SS: 88.6% and UFBS: 58%). As in the case of *postA* and *postB* genes (Matus et al., 2007), *Sce-postC* and *Sce-postD* also share several plesiomorphic features with the ecdysozoan *abdB*, lophotrochozoan *post1* and *post2*, and deuterostome *hox9* – *15* (Fig. S6). These ambiguities, coupled with the lack of chaetognath-specific amino acid motif in the Post genes, complicates the inferences of chaetognath *postA* – *D* evolutionary relationship with bilaterian Post genes. Performing phylogenetic analyses with expanded Hox genes datasets including representatives from Acanthocephala, Micrognathozoa, Gnathostomulida, and other chaetognaths could help identify molecular motifs possibly unique to this enigmatic spiralian lineage and improve the resolution of branching patterns, consequently providing better insights into how these chaetognath posterior class Hox genes relate to those in other bilaterian lineages.

Chaetognaths have retained a complete Hox repertoire, similar to many bilaterians. Our data further support the hypothesis that the last common bilaterian ancestor possessed at least seven Hox genes that comprised anterior, central, and posterior class (Balavoine et al., 2002; De Rosa et al., 1999). The Hox gene complement in chaetognaths seems to have also been shaped by lineage-specific duplications and divergence as evidenced by the presence of *Sce-med6* and *Sce-postA* – *D*, which are likely similar evolutionary events that arose in ecdysozoan *ubx*/*abdA* genes, lophotrochozoan *lox2*/*lox4* genes, and deuterostome *hox9* – *15* genes. In summary, our results demonstrate that chaetognaths exhibit the most extensive repertoire of Hox genes among protostomes, within metazoans only surpassed by chordates.

### 5.3. Staggered Hox expression in Gnathifera

*SceMed4*, a central class Hox gene suggested to belong to PG5, was the first chaetognath Hox gene to have its expression characterized (Papillon et al., 2005). It is expressed as two bilateral clusters in the VNC, providing the earliest evidence that Hox genes may play a role in patterning the ventral CNS of chaetognaths (Papillon et al., 2005). Our data on Hox gene expression in *Spadella* further supports this hypothesis, revealing a staggered expression pattern of 13 Hox genes along the ventral nerve cord in hatchlings (Fig. 3) that persists during the early juvenile stage. This pattern reflects a conserved mechanism observed across many bilaterians, where anterior-posterior (AP) staggered Hox expression is also evident in the neuroectoderm and developing neural structures of particular developmental stages. For instance, staggered Hox gene expression can be inferred in the nervous system of the nemertean *Pantinonemertes californiensis* larva (Hiebert and Maslakova, 2015). It is also observed in the ventral ectoderm, which largely develops into neural structures (e.g. ganglia), during early embryonic stages in several molluscan groups (Barrera Grijalba et al., 2023; Fritsch et al., 2015, 2016; Hinman et al., 2003b; Huan et al., 2020; Samadi and Steiner, 2010; Wollesen et al., 2018). Annelids also typically exhibit a Hox gene expression in a staggered fashion along the AP axis of the nervous system. However, with the exception of the leech *Helobdella* which shows a staggered Hox expression almost exclusively in the developing ventral neural tissues (Kourakis et al., 1997; Nardelli-Haefliger et al., 1994), Hox genes in early development of annelids primarily function in defining overall segment organization and specifying segmental fate rather than being solely involved in patterning the CNS (Bakalenko et al., 2013; Fröbius et al., 2008; Irvine and Martindale, 2000; Kulakova et al., 2007; Wei et al., 2022). In addition, in the rotifer *Brachionus manjavacas* (Fröbius and Funch, 2017) and the brachiopod *Terebratalia transversa* (Schiemann et al., 2017), Hox gene expression is also associated with developing neural structures, although only a partial resemblance of staggered patterning can be observed (Huan et al., 2020). Similarly, a spatially staggered Hox expression pattern is observed, to varying degrees, along the anterior-posterior axis of the developing CNS in arthropods <u>(e.g. Jarvis et al., 2012; Serano et al., 2016; Technau et al., 2006)</u>, hemichordate (Aronowicz and Lowe, 2006), chordates <u>(e.g. Ikuta et al., 2004; Philippidou and Dasen, 2013; Wada et al., 1999)</u>, and acoels (Hejnol and Martindale, 2009; Moreno et al., 2009). In *Drosophila*, Hox genes regulate the identity and specification of region-specific neural subtypes in the CNS (Jarvis et al., 2012). For example, Hox genes contribute to segment-specific patterns of peptidergic neurons, as well as motor neurons, their axonal projections, and target muscle groups (Jarvis et al., 2012; Philippidou and Dasen, 2013; Suska et al., 2011). Similarly, in vertebrates, Hox gene expression in the spinal cord is involved in establishing the columnar identities of motor neurons, with each columnar subtype innervating specific muscle targets along the rostrocaudal (anterior-posterior) axis (Dasen et al., 2003). In *S. cephaloptera*, the staggered Hox expression in the VNC may also contribute to regional specialization (Papillon et al., 2005), potentially resulting in the differentiation of distinct neuronal subtypes along the AP axis. Direct evidence for a spatial correlation between neuronal diversity and Hox gene expression domains in chaetognaths is currently unavailable, but the arrangement of RFamide-immunoreactive (ir) neurons along the AP axis of the VNC offers a potential framework for speculation. These neurons are serially arranged along the AP axis of the VNC, comprising lateral cells (L-group) located anteriorly and dorsal cells (D-group) positioned posteriorly (Goto et al., 1992; Harzsch et al., 2009; Harzsch and Müller, 2007; Harzsch and Wanninger, 2010; Rieger et al., 2011). Conceptually mapping Hox expression domains onto this arrangement, it is plausible that L-group neurons nest within *Sce-hox1* to *Sce-hox3* domains, D1 – D5 align with central-class Hox gene domains, and posterior neurons (D6, D7, and the giant X neuron found only in the Spadellidae) fall within *Sce-medpost* and posterior Hox domains. If such configuration exists, it raises the possibility that Hox genes could influence the diversification and establishment of region-specific of neuronal subtypes in chaetognaths, similar to their functions in *C. elegans* (Zheng et al., 2015), *Drosophila*, and vertebrates (Dasen et al., 2003; Philippidou and Dasen, 2013; Suska et al., 2011). This could point to an ancient and conserved mechanism underlying neural patterning across bilaterians, underscoring further investigation into chaetognath cell type diversity and the molecular processes governing its patterning.

### 5.4. Expression pattern of Hox in non-neural structures of the trunk

Expression domains of several *Spadella* Hox genes are also observed in the mesoderm- and endoderm-derived tissues of the trunk (Fig. 3). *Sce-hox1* has an intense expression around the midtrunk longitudinal muscles which weakens posteriorly, while *Sce-hox8* expression is confined to the longitudinal muscles around the posterior end of the trunk. However, the absence of expression of other Hox genes in the trunk musculature indicates that this is unlikely to reflect a broader role in regionalization along the AP axis. *Hox1*/*labial and hox8/lox2/*lox4 are not commonly implicated in the development of mesodermal structures in spiralians, suggesting that they may have been coopted for the trunk muscle development of *Spadella*. Although further investigation is needed to confirm whether this represents a unique adaptation or an underexplored function in other spiralian lineages.

In contrast, *Sce-hox8*, *Sce-med6*, *Sce-postC*, and *Sce-postD* exhibited spatially coordinated expression in the posterior half of the intestine. Together with *Sce-nk2.1*, the posterior Hox genes expressed in the hindgut may contribute to defining the distribution of light- and dark-granule-containing cells along the intestine. Light-granule cells, which predominate the anterior region and gradually decrease posteriorly, are suggested to be involved in luminal digestive processes, while dark-granule cells, which are most abundant in the posterior region, might play roles in absorption and intracellular digestion (Arnaud et al., 1996; Perez et al., 1999). Additionally, *Sce-postC* and *Sce-postD*, expressed at the posterior tip of the gut, might be involved in specifying rectal cell fate and organizing the formation of the future anus. Hox genes are also sequentially expressed in the developing gut along the AP axis in *Drosophila* (Bienz, 1994), the tunicate *C. intestinalis* (Ikuta et al., 2004), the sea cucumber *Apostichopus japonicus* (Kikuchi et al., 2015), and vertebrates <u>(e.g. Beck et al., 2000; Yokouchi et al., 1995)</u>. In these organisms, posterior Hox genes specifically pattern the posterior regions of the gut. In other bilaterians, Hox gene expression has been observed in specific regions of the digestive tract. For example, *hox6*/*lox5* and *hox3* are expressed in the hindgut of the scaphopod *A. entalis* (Wollesen et al., 2018); *hox1/lab* genes in the foregut-midgut boundary of annelids (Fröbius et al., 2008; Irvine and Martindale, 2000; Kulakova et al., 2007) and in the foregut of the hemichordate *Saccoglossus kowalevskii* (Aronowicz and Lowe, 2006) as well as the amphioxus chordate *Brachiostoma* (Schubert et al., 2005); and *hox3* and *hox7*/*antp* are also observed in the foregut of the polychaete *Capitella* (Fröbius et al., 2008). This distribution of Hox gene expression across different Hox classes in the digestive tract of bilaterians underscores their conserved role in gut regionalization while highlighting lineage-specific adaptations to diverse digestive structures.

There is no observable expression pattern for *Sce-postB*, a duplicate paralog of *Sce-postA*, in the hatchling and early juvenile stages. This absence could be due to several factors. *Sce-postB* expression might be too low or transient to be detected using conventional or HCR in situ hybridization, or the probes designed for its detection may not have worked effectively. It is also plausible that *Sce-postB* is active only at specific developmental stages, such as during embryogenesis or in adults, and may serve specialized functions at these stages. Alternatively, *Sce-postB* could have undergone a partial or complete loss of function, as it is common when one duplicate retains the ancestral role while the other accumulates mutations under relaxed selective pressure (Lynch and Conery, 2000). However, if the lack of detection is due to methodological limitations, it is reasonable to hypothesize that *Sce-postB* might be expressed in similar regions as *Sce-postA*, consistent with the expression patterns of the related duplicates *Sce-postC* and *Sce-postD*.

### 5.5. On the colinearity of chaetognath Hox genes

At present, there is no publicly available genomic data for chaetognaths, and thus there is still no information on the physical linkage of their Hox genes. We generated a draft genome assembly of *S. cephaloptera* (Barrera et al., 2025 – deposited on bioRxiv), but unfortunately, the highly fragmented nature of the assembly prevents the determination of their genomic organization. However, insights can be drawn from rotifers, the closest relatives of chaetognaths with the only available genome data. In the bdelloid rotifer *Adineta vaga*, Hox genes are dispersed across its genome, reflecting extensive genomic rearrangements and reductions, and eventually the loss of the canonical Hox cluster (Flot et al., 2013; Fröbius and Funch, 2017). If chaetognaths share a similar genomic history, it is possible that their Hox genes may not be clustered in the genome, resulting in a non-colinear yet spatially staggered Hox expression pattern, as observed in the tunicates chordates *Oikopleura dioica* (Seo et al., 2004) and *Ciona intestinalis* (Ikuta et al., 2004). Conversely, chaetognaths possess a complete Hox complement unlike rotifers and exhibit staggered expression along the VNC, raising the possibility that their Hox genes could retain some degree of physical linkage similarly to many bilaterians. Nonetheless, generating a more contiguous genome assembly is crucial to determining whether the organization of chaetognath Hox genes, and thus their adherence to spatiotemporal colinearity, aligns with or diverges from those observed in gnathiferans and other spiralians.

### 5.6. Posterior Hox genes pattern the post-anal tail of *Spadella*

*Spadella* exhibits a distinct staggered expression pattern of Hox genes (*Sce-med6*, *Sce-medpost*, *Sce-postA*, *Sce-postC*, and *Sce-postD*) along the tail (Fig. 3), indicating that these posterior Hox genes contribute to the development and regional patterning of the tail. Likewise, posterior-class Hox genes (*hox9 – 13*) in chordates, along with other key gene regulatory network components such as *sonic hedgehog* (*shh*) and *brachyury* (*bra*/*tbxti*), play a critical role in the formation of the post-anal tail (Fodor et al., 2021; Herrmann, 1995; Schwaner et al., 2021). Interestingly, while this suggests potential parallels between the gene networks underlying post-anal tail development, a *shh* ortholog is absent in our *S. cephaloptera* draft transcriptome assembly; although, this does not rule out its presence in the genome. Further, the expression of the *bra* ortholog in the chaetognath *Paraspadella gotoi* is restricted to the stomodeum and blastopore region during early embryonic stages and later localizes around the future mouth in post-gastrulation stages (Takada et al., 2002). This localized expression suggests that *bra* is not involved in tail development. Taken together, these findings suggest that the developmental parallels between chaetognath and chordate post-anal tails represent functional convergence rather than a shared ancestral feature. Consequently, the posterior Hox-mediated patterning observed in the caudal structures of both lineages likely exemplifies a case of convergent evolution, wherein similar developmental genes (i.e., posterior-class Hox genes) independently contribute to tail development.

The chaetognath tail can feature a round-shaped caudal fin, as observed in *Spadella* (Fig. 1a, Fig. S7a), reminiscent of those in some fishes such as soles, and clownfish. In these fishes, this fin shape enables short bursts of propulsion that are crucial for striking prey during ambush or escaping predators (Nursall, 1958). These life strategies align with the behaviors observed in chaetognaths (i.e. feeding and predator response). Thus, the chaetognath tail likely evolved under comparable selective pressures, possibly related to locomotion efficiency, including the adaptation for rapid acceleration during aquatic propulsion. The parallel evolution of post-anal tails in chaetognaths and aquatic chordates underscores how similar ecological demands can drive the emergence of analogous structures through distinct evolutionary pathways.

Tail patterning in chordates represents a canonical function of Hox genes in AP axis regionalization. In contrast, posterior Hox genes in *Spadella* appear to have been recruited for the development of the post-anal tail. Interestingly, some members of Rotifera also possess a post-anal extension, the foot, which serves a similar functional role in locomotion and temporary attachment. Their foot is, however, not patterned by posterior-class Hox genes which are absent in rotifers (Fröbius and Funch, 2017). Instead, in the rotifer *B. manjavacas*, the expression of anterior and central class Hox genes and *medPost* is restricted to neural tissues, including those associated with the foot (Fröbius and Funch, 2017). Thus, the evolutionary trajectories of chaetognaths and rotifers diverged significantly in terms of Hox gene utilization in the development of post-anal structures, suggesting that such adaptations may reflect lineage-specific innovations rather than conserved ancestral roles.

## 6. Conclusion

Our study provides the first comprehensive analysis of anterior (*otx, nk2.1, six3/6*) and Hox gene expression in the chaetognath *Spadella cephaloptera*, shedding light on the molecular mechanisms of anterior-posterior (AP) axis patterning in chaetognaths. We reveal the conserved expression of anterior patterning genes in the cerebral ganglion and head structures, supporting their ancestral role in anterior central nervous system (CNS) specification across bilaterians. Furthermore, we present a staggered expression of Hox genes, including the newly described *Sce-med6*, *Sce-postC*, and *Sce-postD*, along the ventral nerve cord (VNC), hindgut, and post-anal tail. These findings suggest that chaetognaths retain ancestral bilaterian AP patterning mechanisms while exhibiting lineage-specific adaptations. The observed expression patterns resemble neural regionalization patterning in other bilaterians, emphasizing shared developmental mechanisms despite distinct morphologies. As early-diverging spiralians, chaetognaths occupy a key position for understanding spiralian evolution. Their complete Hox complement, in contrast to the reduced Hox repertoire of rotifers, highlights the evolutionary diversity of Hox utilization within Spiralia. Expanding our understanding of AP patterning in chaetognaths will refine our understanding of the ancestral spiralian body plan and the evolutionary emergence of lineage-specific adaptations during spiralian diversification.

## 7. Declarations

### Competing interests

The authors declare that they have no competing interests.

### Funding

This research was funded in whole, or in part, by the Austrian Science Fund (FWF) [P34665].

### Authors’ contributions

JFFO designed and performed the experiments, performed the analyses, and wrote the manuscript. TW designed the experiments, designed and synthesized some riboprobes, secured funding, supervised, and contributed to analyzing the results and to writing of the manuscript. Both authors have read and approved of the final version of the manuscript.

## Acknowledgements

The authors are grateful for financial support by the Faculty of Life Sciences of the University of Vienna.

## Notes

### Competing Interest Statement

The authors have declared no competing interest.

## References

Altschul, S.F., Gish, W., Miller, W., Myers, E.W., Lipman, D.J., 1990. Basic local alignment search tool. J. Mol. Biol. 215, 403–410. 10.1016/S0022-2836(05)80360-2

Andrikou, C., Passamaneck, Y.J., Lowe, C.J., Martindale, M.Q., Hejnol, A., 2019. Molecular patterning during the development of *Phoronopsis harmeri* reveals similarities to rhynchonelliform brachiopods. EvoDevo 10, 33. 10.1186/s13227-019-0146-1

Arenas-Mena, C., Cameron, A.R., Davidson, E.H., 2000. Spatial expression of *Hox* cluster genes in the ontogeny of a sea urchin. Development 127, 4631–4643. 10.1242/dev.127.21.4631

Arnaud, J., Brunet, M., Casanova, J.-P., Mazza, J., Pasqualini, V., 1996. Morphology and ultrastructure of the gut in *Spadella cephaloptera* (Chaetognatha). J. Morphol. 228, 27–44. 10.1002/(SICI)1097-4687(199604)228:1<27::AID-JMOR3>3.0.CO;2-M

Aronowicz, J., Lowe, C.J., 2006. Hox gene expression in the hemichordate *Saccoglossus kowalevskii* and the evolution of deuterostome nervous systems. Integr. Comp. Biol. 46, 890–901. 10.1093/icb/icl045

Bakalenko, N.I., Novikova, E.L., Nesterenko, A.Y., Kulakova, M.A., 2013. Hox gene expression during postlarval development of the polychaete *Alitta virens*. EvoDevo 4, 13. 10.1186/2041-9139-4-13

Balavoine, G., De Rosa, R., Adoutte, A., 2002. Hox clusters and bilaterian phylogeny. Mol. Phylogenet. Evol. 24, 366–373. 10.1016/S1055-7903(02)00237-3

Barrera Grijalba, C.C., Rodríguez Monje, S.V., Gestal, C., Wollesen, T., 2023. Octopod Hox genes and cephalopod plesiomorphies. Sci. Rep. 13, 15492. 10.1038/s41598-023-42435-0

Beck, F., Tata, F., Chawengsaksophak, K., 2000. Homeobox genes and gut development. BioEssays 22, 431–441. 10.1002/(SICI)1521-1878(200005)22:5<431::AID-BIES5>3.0.CO;2-X

Bekkouche, N., Gąsiorowski, L., 2022. Careful amendment of morphological data sets improves phylogenetic frameworks: re-evaluating placement of the fossil *Amiskwia sagittiformis*. J. Syst. Palaeontol. 20, 1–14. 10.1080/14772019.2022.2109217

Bienz, M., 1994. Homeotic genes and positional signalling in the *Drosophila* viscera. Trends Genet. 10, 22–26. 10.1016/0168-9525(94)90015-9

Boyle, M.J., Yamaguchi, E., Seaver, E.C., 2014. Molecular conservation of metazoan gut formation: evidence from expression of endomesoderm genes in *Capitella teleta* (Annelida). EvoDevo 5, 39. 10.1186/2041-9139-5-39

Bruce, A.E.E., Shankland, M., 1998. Expression of the Head Gene Lox22-Otx in the Leech *Helobdella* and the Origin of the Bilaterian Body Plan. Dev. Biol. 201, 101–112. 10.1006/dbio.1998.8968

Carl, M., Loosli, F., Wittbrodt, J., 2002. *Six3* inactivation reveals its essential role for the formation and patterning of the vertebrate eye. Development 129, 4057–4063. 10.1242/dev.129.17.4057

Choi, H.M.T., Beck, V.A., Pierce, N.A., 2014. Next-Generation *in Situ* Hybridization Chain Reaction: Higher Gain, Lower Cost, Greater Durability. ACS Nano 8, 4284–4294. 10.1021/nn405717p

Choi, H.M.T., Calvert, C.R., Husain, N., Huss, D., Barsi, J.C., Deverman, B.E., Hunter, R.C., Kato, M., Lee, S.M., Abelin, A.C.T., Rosenthal, A.Z., Akbari, O.S., Li, Y., Hay, B.A., Sternberg, P.W., Patterson, P.H., Davidson, E.H., Mazmanian, S.K., Prober, D.A., Van De Rijn, M., Leadbetter, J.R., Newman, D.K., Readhead, C., Bronner, M.E., Wold, B., Lansford, R., Sauka-Spengler, T., Fraser, S.E., Pierce, N.A., 2016. Mapping a multiplexed zoo of mRNA expression. Development 143, 3632–3637. 10.1242/dev.140137

Choi, H.M.T., Schwarzkopf, M., Fornace, M.E., Acharya, A., Artavanis, G., Stegmaier, J., Cunha, A., Pierce, N.A., 2018. Third-generation in situ hybridization chain reaction: multiplexed, quantitative, sensitive, versatile, robust. 10.1101/285213

Corbin, J.G., Rutlin, M., Gaiano, N., Fishell, G., 2003. Combinatorial function of the homeodomain proteins Nkx2.1 and Gsh2 in ventral telencephalic patterning. Development 130, 4895–4906. 10.1242/dev.00717

Currie, K.W., Molinaro, A.M., Pearson, B.J., 2016. Neuronal sources of hedgehog modulate neurogenesis in the adult planarian brain. eLife 5, e19735. 10.7554/eLife.19735

Dasen, J.S., Liu, J.-P., Jessell, T.M., 2003. Motor neuron columnar fate imposed by sequential phases of Hox-c activity. Nature 425, 926–933. 10.1038/nature02051

De Rosa, R., Grenier, J.K., Andreeva, T., Cook, C.E., Adoutte, A., Akam, M., Carroll, S.B., Balavoine, G., 1999. Hox genes in brachiopods and priapulids and protostome evolution. Nature 399, 772– 776. 10.1038/21631

Dunn, C.W., Hejnol, A., Matus, D.Q., Pang, K., Browne, W.E., Smith, S.A., Seaver, E., Rouse, G.W., Obst, M., Edgecombe, G.D., Sørensen, M.V., Haddock, S.H.D., Schmidt-Rhaesa, A., Okusu, A., Kristensen, R.M., Wheeler, W.C., Martindale, M.Q., Giribet, G., 2008. Broad phylogenomic sampling improves resolution of the animal tree of life. Nature 452, 745–749. 10.1038/nature06614

Dunn, E.F., Moy, V.N., Angerer, L.M., Angerer, R.C., Morris, R.L., Peterson, K.J., 2007. Molecular paleoecology: using gene regulatory analysis to address the origins of complex life cycles in the late Precambrian. Evol. Dev. 9, 10–24. 10.1111/j.1525-142X.2006.00134.x

Eriksson, B.J., Samadi, L., Schmid, A., 2013. The expression pattern of the genes engrailed, pax6, otd and six3 with special respect to head and eye development in *Euperipatoides kanangrensis* Reid 1996 (Onychophora: Peripatopsidae). Dev. Genes Evol. 223, 237–246. 10.1007/s00427-013-0442-z

Faltine-Gonzalez, D., Havrilak, J., Layden, M.J., 2023. The brain regulatory program predates central nervous system evolution. Sci. Rep. 13, 8626. 10.1038/s41598-023-35721-4

Feuda, R., Peter, I.S., 2022. Homologous gene regulatory networks control development of apical organs and brains in Bilateria. Sci. Adv. 8, eabo2416. 10.1126/sciadv.abo2416

Flot, J.-F., Hespeels, B., Li, X., Noel, B., Arkhipova, I., Danchin, E.G.J., Hejnol, A., Henrissat, B., Koszul, R., Aury, J.-M., Barbe, V., Barthélémy, R.-M., Bast, J., Bazykin, G.A., Chabrol, O., Couloux, A., Da Rocha, M., Da Silva, C., Gladyshev, E., Gouret, P., Hallatschek, O., Hecox-Lea, B., Labadie, K., Lejeune, B., Piskurek, O., Poulain, J., Rodriguez, F., Ryan, J.F., Vakhrusheva, O.A., Wajnberg, E., Wirth, B., Yushenova, I., Kellis, M., Kondrashov, A.S., Mark Welch, D.B., Pontarotti, P., Weissenbach, J., Wincker, P., Jaillon, O., Van Doninck, K., 2013. Genomic evidence for ameiotic evolution in the bdelloid rotifer *Adineta vaga*. Nature 500, 453–457. 10.1038/nature12326

Fodor, A.C.A., Powers, M.M., Andrykovich, K., Liu, J., Lowe, E.K., Brown, C.T., Di Gregorio, A., Stolfi, A., Swalla, B.J., 2021. The Degenerate Tale of Ascidian Tails. Integr. Comp. Biol. 61, 358–369. 10.1093/icb/icab022

Fritsch, M., Wollesen, T., De Oliveira, A.L., Wanninger, A., 2015. Unexpected co-linearity of Hox gene expression in an aculiferan mollusk. BMC Evol. Biol. 15, 151. 10.1186/s12862-015-0414-1

Fritsch, M., Wollesen, T., Wanninger, A., 2016. Hox and ParaHox gene expression in early body plan patterning of polyplacophoran mollusks. J. Exp. Zoolog. B Mol. Dev. Evol. 326, 89–104. 10.1002/jez.b.22671

Fröbius, A.C., Funch, P., 2017. Rotiferan Hox genes give new insights into the evolution of metazoan bodyplans. Nat. Commun. 8, 9. 10.1038/s41467-017-00020-w

Fröbius, A.C., Matus, D.Q., Seaver, E.C., 2008. Genomic organization and expression demonstrate spatial and temporal hox gene colinearity in the lophotrochozoan *Capitella sp. I*. PLoS ONE 3, e4004. 10.1371/journal.pone.0004004

Gąsiorowski, L., Børve, A., Cherneva, I.A., Orús-Alcalde, A., Hejnol, A., 2021. Molecular and morphological analysis of the developing nemertean brain indicates convergent evolution of complex brains in Spiralia. BMC Biol. 19, 175. 10.1186/s12915-021-01113-1

Giribet, G., Distel, D.L., Polz, M., Sterrer, W., Wheeler, W.C., 2000. Triploblastic Relationships with Emphasis on the Acoelomates and the Position of Gnathostomulida, Cycliophora, Plathelminthes, and Chaetognatha: A Combined Approach of 18S rDNA Sequences and Morphology. Syst. Biol. 49, 539–562. 10.1080/10635159950127385

Gonzalez, P., Uhlinger, K.R., Lowe, C.J., 2017. The Adult Body Plan of Indirect Developing Hemichordates Develops by Adding a Hox-Patterned Trunk to an Anterior Larval Territory. Curr. Biol. 27, 87–95. 10.1016/j.cub.2016.10.047

Goto, T., Katayama-Kumoi, Y., Tohyama, M., Yoshida, M., 1992. Distribution and development of the serotonin-and RFamide-like immunoreactive neurons in the arrowworm, *Paraspadella gotoi* (Chaetognatha). Cell Tissue Res. 267, 215–222. 10.1007/BF00302958

Guindon, S., Dufayard, J.-F., Lefort, V., Anisimova, M., Hordijk, W., Gascuel, O., 2010. New Algorithms and Methods to Estimate Maximum-Likelihood Phylogenies: Assessing the Performance of PhyML 3.0. Syst. Biol. 59, 307–321. 10.1093/sysbio/syq010

Harfe, B.D., Fire, A., 1998. Muscle and nerve-specific regulation of a novel NK-2 class homeodomain factor in *Caenorhabditis elegans*. Development 125, 421–429. 10.1242/dev.125.3.421

Harzsch, S., Müller, C.H., 2007. A new look at the ventral nerve centre of *Sagitta*: implications for the phylogenetic position of Chaetognatha (arrow worms) and the evolution of the bilaterian nervous system. Front. Zool. 4, 14. 10.1186/1742-9994-4-14

Harzsch, S., Müller, C.H.G., Rieger, V., Perez, Y., Sintoni, S., Sardet, C., Hansson, B., 2009. Fine structure of the ventral nerve centre and interspecific identification of individual neurons in the enigmatic Chaetognatha. Zoomorphology 128, 53–73. 10.1007/s00435-008-0074-4

Harzsch, S., Wanninger, A., 2010. Evolution of invertebrate nervous systems: the Chaetognatha as a case study. Acta Zool. 91, 35–43. 10.1111/j.1463-6395.2009.00423.x

Hejnol, A., Lowe, C.J., 2015. Embracing the comparative approach: how robust phylogenies and broader developmental sampling impacts the understanding of nervous system evolution. Philos. Trans. R. Soc. Lond. B. Biol. Sci. 370, 20150045. 10.1098/rstb.2015.0045

Hejnol, A., Martindale, M.Q., 2009. Coordinated spatial and temporal expression of Hox genes during embryogenesis in the acoel *Convolutriloba longifissura*. BMC Biol. 7, 65. 10.1186/1741-7007-7-65

Hejnol, A., Martindale, M.Q., 2008. Acoel development indicates the independent evolution of the bilaterian mouth and anus. Nature 456, 382–386. 10.1038/nature07309

Helfenbein, K.G., Fourcade, H.M., Vanjani, R.G., Boore, J.L., 2004. The mitochondrial genome of *Paraspadella gotoi* is highly reduced and reveals that chaetognaths are a sister group to protostomes. Proc. Natl. Acad. Sci. 101, 10639–10643. 10.1073/pnas.0400941101

Herrmann, B.G., 1995. The mouse Brachyury (T) gene. Semin. Dev. Biol. 6, 385–394. 10.1016/S1044-5781(06)80002-2

Hiebert, L.S., Maslakova, S.A., 2015. Expression of Hox, Cdx, and Six3/6 genes in the hoplonemertean *Pantinonemertes californiensis* offers insight into the evolution of maximally indirect development in the phylum Nemertea. EvoDevo 6, 26. 10.1186/s13227-015-0021-7

Hinman, V.F., Nguyen, A.T., Davidson, E.H., 2003a. Expression and function of a starfish Otx ortholog, AmOtx: a conserved role for Otx proteins in endoderm development that predates divergence of the eleutherozoa. Mech. Dev. 120, 1165–1176. 10.1016/j.mod.2003.08.002

Hinman, V.F., O’Brien, E.K., Richards, G.S., Degnan, B.M., 2003b. Expression of anterior *Hox* genes during larval development of the gastropod *Haliotis asinina*. Evol. Dev. 5, 508–521. 10.1046/j.1525-142X.2003.03056.x

Hoang, D.T., Chernomor, O., Von Haeseler, A., Minh, B.Q., Vinh, L.S., 2018. UFBoot2: Improving the Ultrafast Bootstrap Approximation. Mol. Biol. Evol. 35, 518–522. 10.1093/molbev/msx281

Holland, L.Z., Carvalho, J.E., Escriva, H., Laudet, V., Schubert, M., Shimeld, S.M., Yu, J.-K., 2013. Evolution of bilaterian central nervous systems: a single origin? EvoDevo 4, 27. 10.1186/2041-9139-4-27

Huan, P., Wang, Q., Tan, S., Liu, B., 2020. Dorsoventral decoupling of Hox gene expression underpins the diversification of molluscs. Proc. Natl. Acad. Sci. 117, 503–512. 10.1073/pnas.1907328117

Ikuta, T., Yoshida, N., Satoh, N., Saiga, H., 2004. *Ciona intestinalis* Hox gene cluster: Its dispersed structure and residual colinear expression in development. Proc. Natl. Acad. Sci. 101, 15118– 15123. 10.1073/pnas.0401389101

Irvine, S.Q., Martindale, M.Q., 2000. Expression Patterns of Anterior Hox Genes in the Polychaete *Chaetopterus*: Correlation with Morphological Boundaries. Dev. Biol. 217, 333–351. 10.1006/dbio.1999.9541

Janssen, R., Eriksson, B., Tait, N.N., Budd, G.E., 2014. Onychophoran Hox genes and the evolution of arthropod Hox gene expression. Front. Zool. 11, 22. 10.1186/1742-9994-11-22

Jarvis, E., Bruce, H.S., Patel, N.H., 2012. Evolving specialization of the arthropod nervous system. Proc. Natl. Acad. Sci. 109, 10634–10639. 10.1073/pnas.1201876109

Kalyaanamoorthy, S., Minh, B.Q., Wong, T.K.F., Von Haeseler, A., Jermiin, L.S., 2017. ModelFinder: fast model selection for accurate phylogenetic estimates. Nat. Methods 14, 587–589. 10.1038/nmeth.4285

Katoh, K., 2005. MAFFT version 5: improvement in accuracy of multiple sequence alignment. Nucleic Acids Res. 33, 511–518. 10.1093/nar/gki198

Katoh, K., 2002. MAFFT: a novel method for rapid multiple sequence alignment based on fast Fourier transform. Nucleic Acids Res. 30, 3059–3066. 10.1093/nar/gkf436

Kaul-Strehlow, S., Urata, M., Praher, D., Wanninger, A., 2017. Neuronal patterning of the tubular collar cord is highly conserved among enteropneusts but dissimilar to the chordate neural tube. Sci. Rep. 7, 7003. 10.1038/s41598-017-07052-8

Kerbl, A., Martín-Durán, J.M., Worsaae, K., Hejnol, A., 2016. Molecular regionalization in the compact brain of the meiofaunal annelid *Dinophilus gyrociliatus* (Dinophilidae). EvoDevo 7, 20. 10.1186/s13227-016-0058-2

Kikuchi, M., Omori, A., Kurokawa, D., Akasaka, K., 2015. Patterning of anteroposterior body axis displayed in the expression of Hox genes in sea cucumber *Apostichopus japonicus*. Dev. Genes Evol. 225, 275–286. 10.1007/s00427-015-0510-7

Klann, M., Seaver, E.C., 2019. Functional role of pax6 during eye and nervous system development in the annelid *Capitella teleta*. Dev. Biol. 456, 86–103. 10.1016/j.ydbio.2019.08.011

Kourakis, M.J., Master, V.A., Lokhorst, D.K., Nardelli-Haefliger, D., Wedeen, C.J., Martindale, M.Q., Shankland, M., 1997. Conserved Anterior Boundaries of Hox Gene Expression in the Central Nervous System of the Leech *Helobdella*. Dev. Biol. 190, 284–300. 10.1006/dbio.1997.8689

Kuehn, E., Clausen, D.S., Null, R.W., Metzger, B.M., Willis, A.D., Özpolat, B.D., 2022. Segment number threshold determines juvenile onset of germline cluster expansion in *Platynereis dumerilii*. J. Exp. Zoolog. B Mol. Dev. Evol. 338, 225–240. 10.1002/jez.b.23100

Kulakova, M., Bakalenko, N., Novikova, E., Cook, C.E., Eliseeva, E., Steinmetz, P.R.H., Kostyuchenko, R.P., Dondua, A., Arendt, D., Akam, M., Andreeva, T., 2007. Hox gene expression in larval development of the polychaetes *Nereis virens* and *Platynereis dumerilii* (Annelida, Lophotrochozoa). Dev. Genes Evol. 217, 39–54. 10.1007/s00427-006-0119-y

Laumer, C.E., Fernández, R., Lemer, S., Combosch, D., Kocot, K.M., Riesgo, A., Andrade, S.C.S., Sterrer, W., Sørensen, M.V., Giribet, G., 2019. Revisiting metazoan phylogeny with genomic sampling of all phyla. Proc. R. Soc. B Biol. Sci. 286, 20190831. 10.1098/rspb.2019.0831

Littlewood, D.T.J., Telford, M.J., Clough, K.A., Rohde, K., 1998. Gnathostomulida—An Enigmatic Metazoan Phylum from both Morphological and Molecular Perspectives. Mol. Phylogenet. Evol. 9, 72–79. 10.1006/mpev.1997.0448

Loosli, F., Köster, R.W., Carl, M., Krone, A., Wittbrodt, J., 1998. Six3, a medaka homologue of the *Drosophila* homeobox gene sine oculis is expressed in the anterior embryonic shield and the developing eye. Mech. Dev. 74, 159–164. 10.1016/S0925-4773(98)00055-0

Lowe, C.J., Wu, M., Salic, A., Evans, L., Lander, E., Stange-Thomann, N., Gruber, C.E., Gerhart, J., Kirschner, M., 2003. Anteroposterior Patterning in Hemichordates and the Origins of the Chordate Nervous System. Cell 113, 853–865. 10.1016/S0092-8674(03)00469-0

Luo, Y.-J., Kanda, M., Koyanagi, R., Hisata, K., Akiyama, T., Sakamoto, H., Sakamoto, T., Satoh, N., 2017. Nemertean and phoronid genomes reveal lophotrochozoan evolution and the origin of bilaterian heads. Nat. Ecol. Evol. 2, 141–151. 10.1038/s41559-017-0389-y

Lynch, M., Conery, J.S., 2000. The Evolutionary Fate and Consequences of Duplicate Genes. Science 290, 1151–1155. 10.1126/science.290.5494.1151

Mallatt, J., Winchell, C.J., 2002. Testing the New Animal Phylogeny: First Use of Combined Large-Subunit and Small-Subunit rRNA Gene Sequences to Classify the Protostomes. Mol. Biol. Evol. 19, 289–301. 10.1093/oxfordjournals.molbev.a004082

Marlétaz, F., Martin, E., Perez, Y., Papillon, D., Caubit, X., Lowe, C.J., Freeman, B., Fasano, L., Dossat, C., Wincker, P., Weissenbach, J., Le Parco, Y., 2006. Chaetognath phylogenomics: a protostome with deuterostome-like development. Curr. Biol. 16, R577–R578. 10.1016/j.cub.2006.07.016

Marlétaz, F., Peijnenburg, K.T.C.A., Goto, T., Satoh, N., Rokhsar, D.S., 2019. A New Spiralian Phylogeny Places the Enigmatic Arrow Worms among Gnathiferans. Curr. Biol. 29, 312–318.e3. 10.1016/j.cub.2018.11.042

Marlow, H., Tosches, M.A., Tomer, R., Steinmetz, P.R., Lauri, A., Larsson, T., Arendt, D., 2014. Larval body patterning and apical organs are conserved in animal evolution. BMC Biol. 12, 7. 10.1186/1741-7007-12-7

Martín-Durán, J.M., Pang, K., Børve, A., Lê, H.S., Furu, A., Cannon, J.T., Jondelius, U., Hejnol, A., 2018. Convergent evolution of bilaterian nerve cords. Nature 553, 45–50. 10.1038/nature25030

Martín-Durán, J.M., Vellutini, B.C., Hejnol, A., 2015. Evolution and development of the adelphophagic, intracapsular Schmidt’s larva of the nemertean *Lineus ruber*. EvoDevo 6, 28. 10.1186/s13227-015-0023-5

Matus, D.Q., Copley, R.R., Dunn, C.W., Hejnol, A., Eccleston, H., Halanych, K.M., Martindale, M.Q., Telford, M.J., 2006. Broad taxon and gene sampling indicate that chaetognaths are protostomes. Curr. Biol. 16, R575–R576. 10.1016/j.cub.2006.07.017

Matus, D.Q., Halanych, K.M., Martindale, M.Q., 2007. The Hox gene complement of a pelagic chaetognath, *Flaccisagitta enflata*. Integr. Comp. Biol. 47, 854–864. 10.1093/icb/icm077

Minh, B.Q., Schmidt, H.A., Chernomor, O., Schrempf, D., Woodhams, M.D., Von Haeseler, A., Lanfear, R., 2020. IQ-TREE 2: New Models and Efficient Methods for Phylogenetic Inference in the Genomic Era. Mol. Biol. Evol. 37, 1530–1534. 10.1093/molbev/msaa015

Moreno, E., Nadal, M., Baguñà, J., Martínez, P., 2009. Tracking the origins of the bilaterian *Hox* patterning system: insights from the acoel flatworm *Symsagittifera roscoffensis*. Evol. Dev. 11, 574–581. 10.1111/j.1525-142X.2009.00363.x

Murakami, Y., Ogasawara, M., Sugahara, F., Hirano, S., Satoh, N., Kuratani, S., 2001. Identification and expression of the lamprey *Pax6* gene: evolutionary origin of the segmented brain of vertebrates. Development 128, 3521–3531. 10.1242/dev.128.18.3521

Nardelli-Haefliger, D., Bruce, A.E.E., Shankland, M., 1994. An axial domain of HOM/Hox gene expression is formed by morphogenetic alignment of independently specified cell lineages in the leech *Helobdella*. Development 120, 1839–1849. 10.1242/dev.120.7.1839

Nursall, J.R., 1958. The Caudal Fin as a Hydrofoil. Evolution 12, 116. 10.2307/2405913

Ogura, A., Yoshida, M., Moritaki, T., Okuda, Y., Sese, J., Shimizu, K.K., Sousounis, K., Tsonis, P.A., 2013. Loss of the six3/6 controlling pathways might have resulted in pinhole-eye evolution in Nautilus. Sci. Rep. 3, 1432. 10.1038/srep01432

Oliver, G., Mailhos, A., Wehr, R., Copeland, N.G., Jenkins, N.A., Gruss, P., 1995. *Six3*, a murine homologue of the *sine oculis* gene, demarcates the most anterior border of the developing neural plate and is expressed during eye development. Development 121, 4045–4055. 10.1242/dev.121.12.4045

Omori, A., Akasaka, K., Kurokawa, D., Amemiya, S., 2011. Gene expression analysis of Six3, Pax6, and Otx in the early development of the stalked crinoid *Metacrinus rotundus*. Gene Expr. Patterns 11, 48–56. 10.1016/j.gep.2010.09.002

Ordoñez, J.F., Wollesen, T., 2024. Unfolding the ventral nerve center of chaetognaths. Neural Develop. 19, 5. 10.1186/s13064-024-00182-6

Papillon, D., Perez, Y., Caubit, X., Le Parco, Y., 2004. Identification of chaetognaths as protostomes ps supported by the analysis of their mitochondrial genome. Mol. Biol. Evol. 21, 2122–2129. 10.1093/molbev/msh229

Papillon, D., Perez, Y., Fasano, L., Le Parco, Y., Caubit, X., 2005. Restricted expression of a median Hox gene in the central nervous system of chaetognaths. Dev. Genes Evol. 215, 369–373. 10.1007/s00427-005-0483-z

Papillon, D., Perez, Y., Fasano, L., Le Parco, Y., Caubit, X., 2003. Hox gene survey in the chaetognath *Spadella cephaloptera*: evolutionary implications. Dev. Genes Evol. 213, 142–148. 10.1007/s00427-003-0306-z

Paps, J., Baguñà, J., Riutort, M., 2009. Lophotrochozoa internal phylogeny: new insights from an up-to-date analysis of nuclear ribosomal genes. Proc. R. Soc. B Biol. Sci. 276, 1245–1254. 10.1098/rspb.2008.1574

Park, T.-Y.S., Nielsen, M.L., Parry, L.A., Sørensen, M.V., Lee, M., Kihm, J.-H., Ahn, I., Park, C., De Vivo, G., Smith, M.P., Harper, D.A.T., Nielsen, A.T., Vinther, J., 2024. A giant stem-group chaetognath. Sci. Adv. 10, eadi6678. 10.1126/sciadv.adi6678

Perez, Y., Arnaud, J., Brunet, M., Casanova, J.-P., Mazza, J., 1999. Morphological study of the gut in *Sagitta setosa*, *S. serratodentata* and *S. pacifica* (Chaetognatha). Functional implications in digestive processes. J. Mar. Biol. Assoc. U. K. 79, 1097–1109. 10.1017/S0025315499001356

Perry, K.J., Lyons, D.C., Truchado-Garcia, M., Fischer, A.H.L., Helfrich, L.W., Johansson, K.B., Diamond, J.C., Grande, C., Henry, J.Q., 2015. Deployment of regulatory genes during gastrulation and germ layer specification in a model spiralian mollusc *Crepidula*. Dev. Dyn. 244, 1215–1248. 10.1002/dvdy.24308

Peterson, K.J., Eernisse, D.J., 2001. Animal phylogeny and the ancestry of bilaterians: inferences from morphology and 18S rDNA gene sequences. Evol. Dev. 3, 170–205. 10.1046/j.1525-142x.2001.003003170.x

Philippe, H., Brinkmann, H., Copley, R.R., Moroz, L.L., Nakano, H., Poustka, A.J., Wallberg, A., Peterson, K.J., Telford, M.J., 2011. Acoelomorph flatworms are deuterostomes related to *Xenoturbella*. Nature 470, 255–258. 10.1038/nature09676

Philippidou, P., Dasen, J.S., 2013. Hox Genes: Choreographers in Neural Development, Architects of Circuit Organization. Neuron 80, 12–34. 10.1016/j.neuron.2013.09.020

Redl, E., Scherholz, M., Wollesen, T., Todt, C., Wanninger, A., 2018. Expression of six3 and otx in Solenogastres (Mollusca) supports an ancestral role in bilaterian anterior-posterior axis patterning. Evol. Dev. 20, 17–28. 10.1111/ede.12245

Rieger, V., Perez, Y., Müller, C.H.G., Lacalli, T., Hansson, B.S., Harzsch, S., 2011. Development of the nervous system in hatchlings of *Spadella cephaloptera* (Chaetognatha), and implications for nervous system evolution in Bilateria: Nervous system development in S. cephaloptera. Dev. Growth Differ. 53, 740–759. 10.1111/j.1440-169X.2011.01283.x

Ristoratore, F., Spagnuolo, A., Aniello, F., Branno, M., Fabbrini, F., Lauro, R.D., 1999. Expression and functional analysis of *Cititf1*, an ascidian *NK-2* class gene, suggest its role in endoderm development. Development 126, 5149–5159. 10.1242/dev.126.22.5149

Samadi, L., Steiner, G., 2010. Expression of Hox genes during the larval development of the snail, *Gibbula varia* (L.)—further evidence of non-colinearity in molluscs. Dev. Genes Evol. 220, 161– 172. 10.1007/s00427-010-0338-0

Santagata, S., Resh, C., Hejnol, A., Martindale, M.Q., Passamaneck, Y.J., 2012. Development of the larval anterior neurogenic domains of *Terebratalia transversa* (Brachiopoda) provides insights into the diversification of larval apical organs and the spiralian nervous system. EvoDevo 3, 3. 10.1186/2041-9139-3-3

Schiemann, S.M., Martín-Durán, J.M., Børve, A., Vellutini, B.C., Passamaneck, Y.J., Hejnol, A., 2017. Clustered brachiopod Hox genes are not expressed collinearly and are associated with lophotrochozoan novelties. Proc. Natl. Acad. Sci. 114. 10.1073/pnas.1614501114

Schindelin, J., Arganda-Carreras, I., Frise, E., Kaynig, V., Longair, M., Pietzsch, T., Preibisch, S., Rueden, C., Saalfeld, S., Schmid, B., Tinevez, J.Y., White, D.J., Hartenstein, V., Eliceiri, K., Tomancak, P., Cardona, A., 2012. Fiji: An open-source platform for biological-image analysis. Nat. Methods. 10.1038/nmeth.2019

Schubert, M., Yu, J.-K., Holland, N.D., Escriva, H., Laudet, V., Holland, L.Z., 2005. Retinoic acid signaling acts via Hox1 to establish the posterior limit of the pharynx in the chordate amphioxus. Development 132, 61–73. 10.1242/dev.01554

Schwaner, M.J., Hsieh, S.T., Braasch, I., Bradley, S., Campos, C.B., Collins, C.E., Donatelli, C.M., Fish, F.E., Fitch, O.E., Flammang, B.E., Jackson, B.E., Jusufi, A., Mekdara, P.J., Patel, A., Swalla, B.J., Vickaryous, M., McGowan, C.P., 2021. Future Tail Tales: A Forward-Looking, Integrative Perspective on Tail Research. Integr. Comp. Biol. 61, 521–537. 10.1093/icb/icab082

Scimone, M.L., Kravarik, K.M., Lapan, S.W., Reddien, P.W., 2014. Neoblast specialization in regeneration of the planarian *Schmidtea mediterranea*. Stem Cell Rep. 3, 339–352. 10.1016/j.stemcr.2014.06.001

Seo, H.-C., Edvardsen, R.B., Maeland, A.D., Bjordal, M., Jensen, M.F., Hansen, A., Flaat, M., Weissenbach, J., Lehrach, H., Wincker, P., Reinhardt, R., Chourrout, D., 2004. Hox cluster disintegration with persistent anteroposterior order of expression in *Oikopleura dioica*. Nature 431, 67–71. 10.1038/nature02709

Serano, J.M., Martin, A., Liubicich, D.M., Jarvis, E., Bruce, H.S., La, K., Browne, W.E., Grimwood, J., Patel, N.H., 2016. Comprehensive analysis of Hox gene expression in the amphipod crustacean *Parhyale hawaiensis*. Dev. Biol. 409, 297–309. 10.1016/j.ydbio.2015.10.029

Steenwyk, J.L., Buida, T.J., Li, Y., Shen, X.-X., Rokas, A., 2020. ClipKIT: A multiple sequence alignment trimming software for accurate phylogenomic inference. PLOS Biol. 18, e3001007. 10.1371/journal.pbio.3001007

Steinmetz, P.R., Urbach, R., Posnien, N., Eriksson, J., Kostyuchenko, R.P., Brena, C., Guy, K., Akam, M., Bucher, G., Arendt, D., 2010. Six3 demarcates the anterior-most developing brain region in bilaterian animals. EvoDevo 1, 14. 10.1186/2041-9139-1-14

Steinmetz, P.R.H., Kostyuchenko, R.P., Fischer, A., Arendt, D., 2011. The segmental pattern of otx, gbx, and Hox genes in the annelid *Platynereis dumerilii*: The segmental pattern of otx, gbx, and Hox genes. Evol. Dev. 13, 72–79. 10.1111/j.1525-142X.2010.00457.x

Suska, A., Miguel-Aliaga, I., Thor, S., 2011. Segment-specific generation of *Drosophila* Capability neuropeptide neurons by multi-faceted Hox cues. Dev. Biol. 353, 72–80. 10.1016/j.ydbio.2011.02.015

Takacs, C.M., Moy, V.N., Peterson, K.J., 2002. Testing putative hemichordate homologues of the chordate dorsal nervous system and endostyle: expression of *NK2.1* (*TTF-1*) in the acorn worm *Ptychodera flava* (Hemichordata, Ptychoderidae). Evol. Dev. 4, 405–417. 10.1046/j.1525-142X.2002.02029.x

Takada, N., Goto, T., Satoh, N., 2002. Expression pattern of the *Brachyury* gene in the arrow worm *Paraspadella gotoi* (chaetognatha). genesis 32, 240–245. 10.1002/gene.10077

Technau, G.M., Berger, C., Urbach, R., 2006. Generation of cell diversity and segmental pattern in the embryonic central nervous system of *Drosophila*. Dev. Dyn. 235, 861–869. 10.1002/dvdy.20566

Telford, M.J., Holland, P.W., 1993. The phylogenetic affinities of the chaetognaths: a molecular analysis. Mol. Biol. Evol. 10, 660–76. 10.1093/oxfordjournals.molbev.a040030

Tessmar-Raible, K., Raible, F., Christodoulou, F., Guy, K., Rembold, M., Hausen, H., Arendt, D., 2007. Conserved sensory-neurosecretory cell types in annelid and fish forebrain: Insights into hypothalamus evolution. Cell 129, 1389–1400. 10.1016/j.cell.2007.04.041

Tsuneoka, Y., Funato, H., 2020. Modified in situ Hybridization Chain Reaction Using Short Hairpin DNAs. Front. Mol. Neurosci. 13, 75. 10.3389/fnmol.2020.00075

Venkatesh, T.V., Holland, N.D., Holland, L.Z., Su, M.-T., Bodmer, R., 1999. Sequence and developmental expression of amphioxus AmphiNk2-1: insights into the evolutionary origin of the vertebrate thyroid gland and forebrain. Dev. Genes Evol. 209, 254–259. 10.1007/s004270050250

Vöcking, O., Kourtesis, I., Hausen, H., 2015. Posterior eyespots in larval chitons have a molecular identity similar to anterior cerebral eyes in other bilaterians. EvoDevo 6, 40. 10.1186/s13227-015-0036-0

Wada, H., Garcia-Fernàndez, J., Holland, P.W.H., 1999. Colinear and Segmental Expression of Amphioxus Hox Genes. Dev. Biol. 213, 131–141. 10.1006/dbio.1999.9369

Wang, Y., Liu, X., Zeng, Y., Saka, S.K., Xie, W., Goldaracena, I., Kohman, R.E., Yin, P., Church, G.M., 2024. Multiplexed *in situ* protein imaging using DNA-barcoded antibodies with extended hybridization chain reactions. Nucleic Acids Res. gkae592. 10.1093/nar/gkae592

Wei, M., Qin, Z., Kong, D., Liu, D., Zheng, Q., Bai, S., Zhang, Z., Ma, Y., 2022. Echiuran Hox genes provide new insights into the correspondence between Hox subcluster organization and collinearity pattern. Proc. R. Soc. B Biol. Sci. 289, 20220705. 10.1098/rspb.2022.0705

Wollesen, T., Rodríguez Monje, S.V., Luiz De Oliveira, A., Wanninger, A., 2018. Staggered Hox expression is more widespread among molluscs than previously appreciated. Proc. R. Soc. B Biol. Sci. 285, 20181513. 10.1098/rspb.2018.1513

Wollesen, T., Rodriguez Monje, S. V, Oel, A.P., Arendt, D., 2023. Characterization of eyes, photoreceptors, and opsins in developmental stages of the arrow worm *Spadella cephaloptera* (Chaetognatha). J. Exp. Zoolog. B Mol. Dev. Evol. 10.1002/jez.b.23193

Wollesen, T., Scherholz, M., Rodríguez Monje, S.V., Redl, E., Todt, C., Wanninger, A., 2017. Brain regionalization genes are co-opted into shell field patterning in Mollusca. Sci. Rep. 7, 5486. 10.1038/s41598-017-05605-5

Yokouchi, Y., Sakiyama, J.-I., Kuroiwa, A., 1995. Coordinated expression of Abd-B subfamily genes of the HoxA cluster in the developing digestive tract of chick embryo. Dev. Biol. 169, 76–89. 10.1006/dbio.1995.1128

Zaffran, S., Das, G., Frasch, M., 2000. The NK-2 homeobox gene scarecrow (scro) is expressed in pharynx, ventral nerve cord and brain of *Drosophila* embryos. Mech. Dev. 94, 237–241. 10.1016/S0925-4773(00)00298-7

Zheng, C., Diaz-Cuadros, M., Chalfie, M., 2015. Hox Genes Promote Neuronal Subtype Diversification through Posterior Induction in *Caenorhabditis elegans*. Neuron 88, 514–527. 10.1016/j.neuron.2015.09.049

