## Supplementary Figures for "Chaetognaths exhibit the most extensive repertoire of Hox genes among protostomes"

Tim Wollesen

June F. Ordoñez

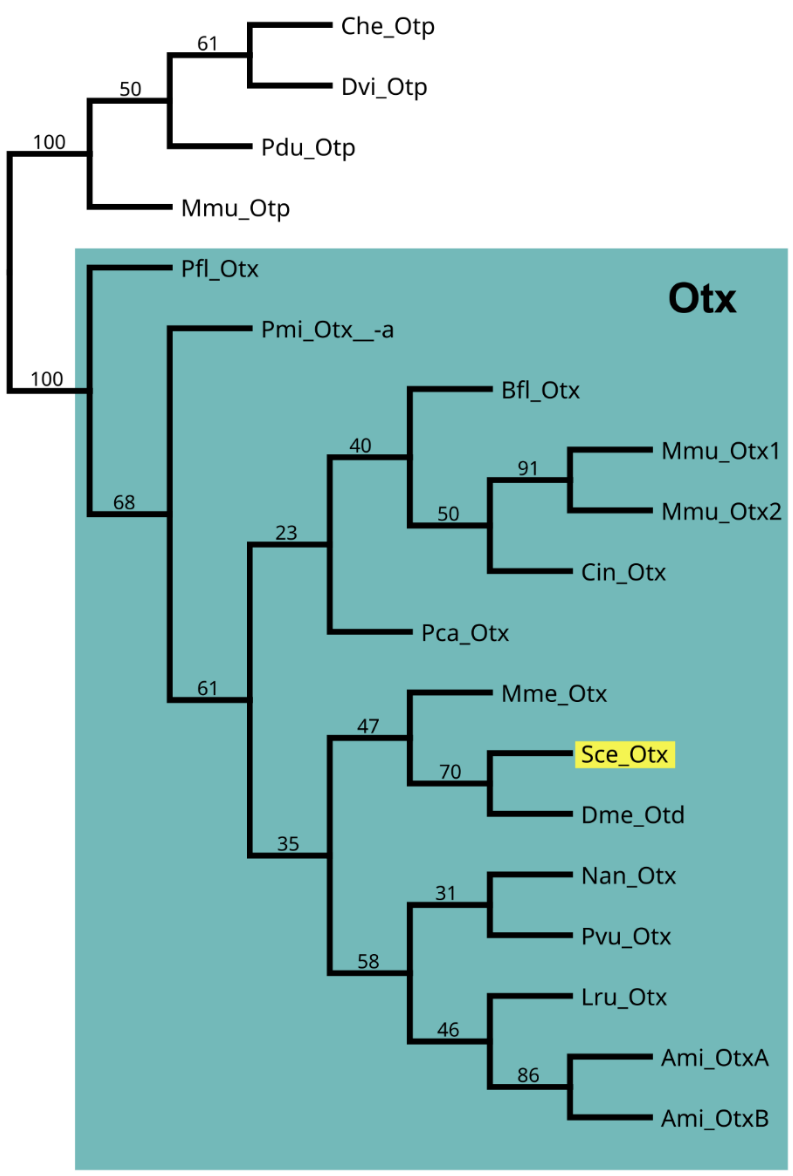


**Figure S1.** Orthology analysis of the *otx* transcript of *Spadella cephaloptera***.** The phylogenetic tree of *otx* genes is based on bilaterian protein sequences obtained from published literature and BLAST searches of the NCBI GenBank. The tree was generated using Maximum Likelihood analysis implemented in IQTREE with the following configurations: VT + G was selected based on ModelFinder and ultrafast bootstrap set to 1000. The support values of branches indicate maximum likelihood bootstrap values. The tree is rooted with *otp* (*orthopedia*). The otx group is highlighted in the teal box and *Sce-otx* in yellow. Species abbreviations are in Table S1.

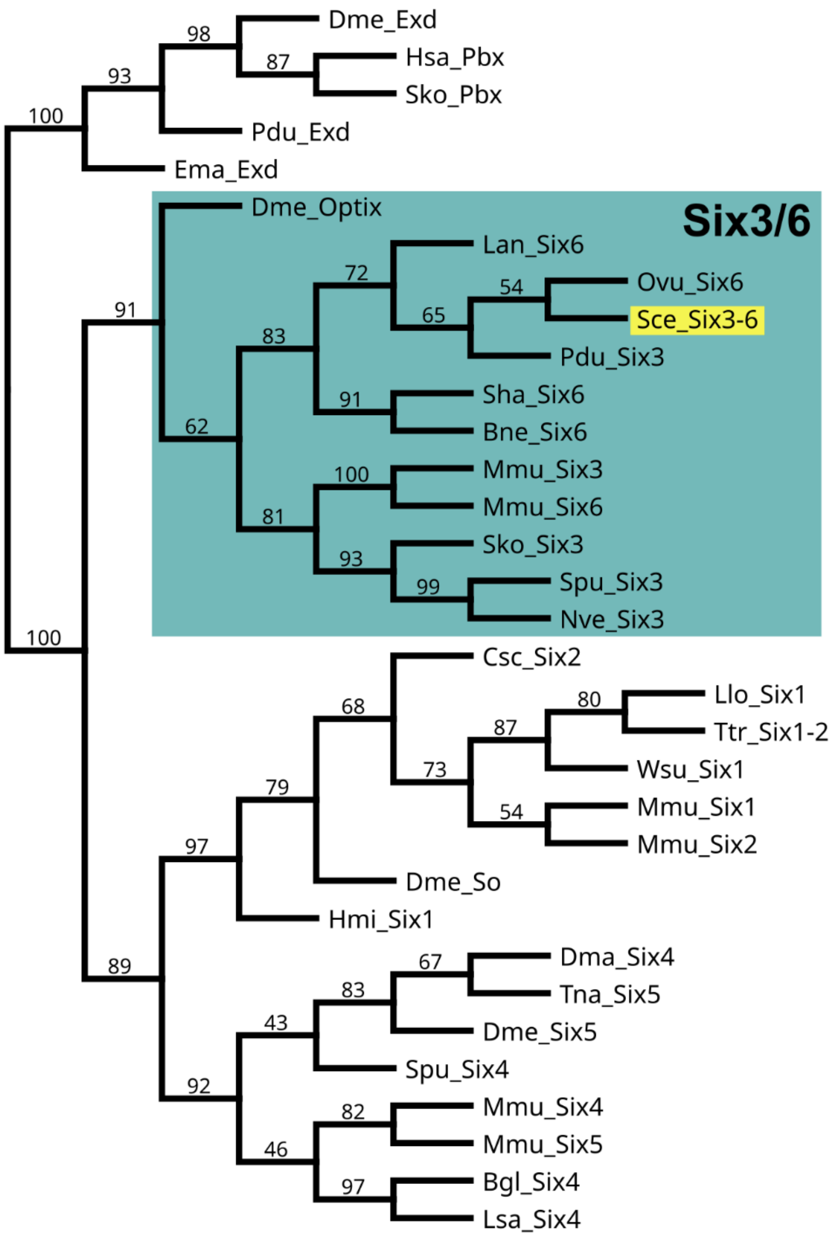


**Figure S2.** Orthology analysis of the *six3/6* transcript of *Spadella cephaloptera***.** The phylogenetic tree of *six* genes is based on bilaterian protein sequences obtained from published literature and BLAST searches of the NCBI GenBank. The tree was generated using Maximum Likelihood analysis implemented in IQTREE with the following configurations: LG + G was selected based on ModelFinder and ultrafast bootstrap set to 1000. The support values of branches indicate maximum likelihood bootstrap values. The tree is rooted with *extradenticle* (*exd*)/ pre-B-cell leukemia homeobox 1 (*pbx*) as an outgroup. The six3/6 group is highlighted in the teal box and *Sce-six3/6* in yellow. Species abbreviations are in Table S1.

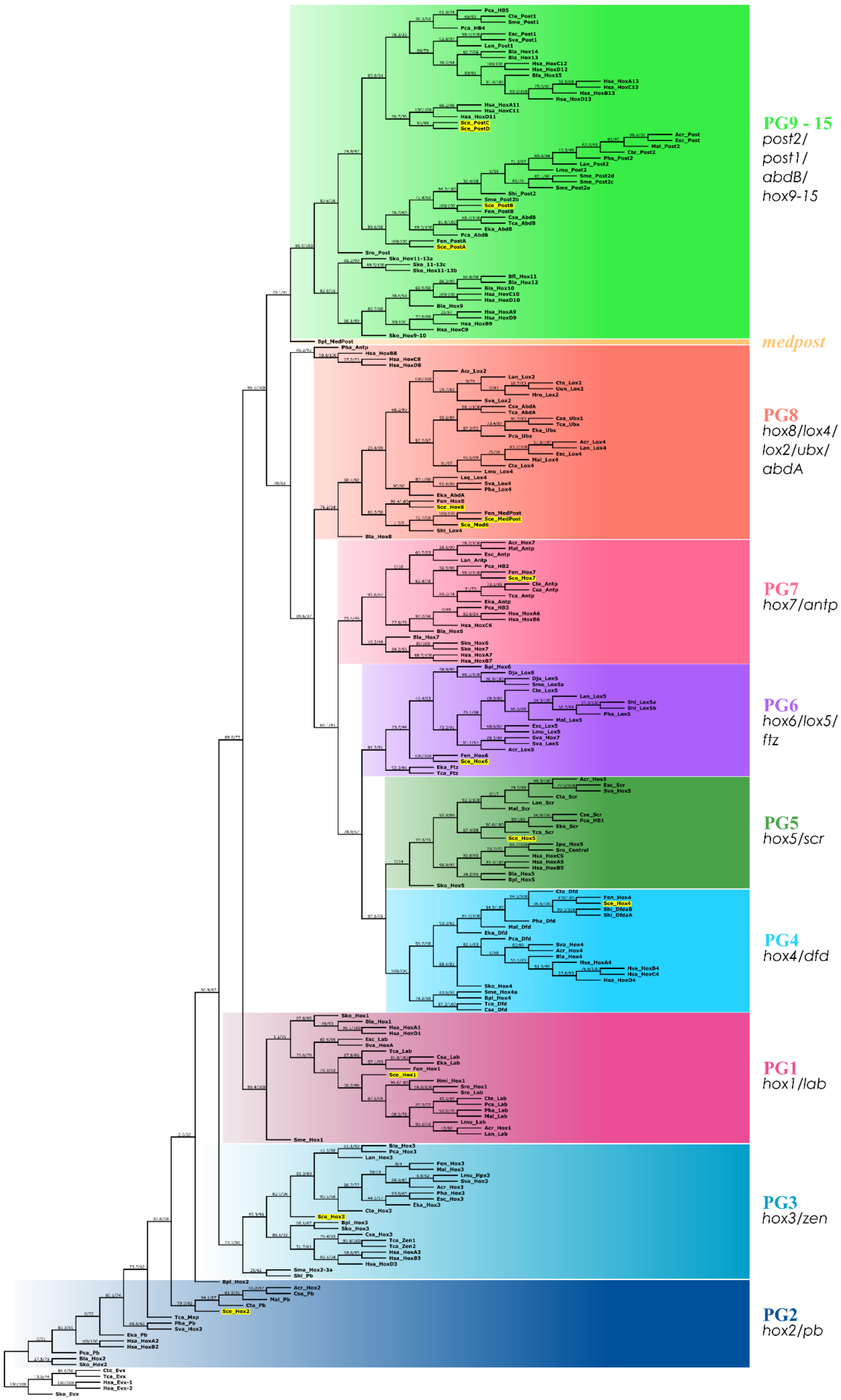

PG9 - 15
post2/
post1/
abdB/
hox9-15
medpost
PG8
hox8/lox4/
lox2/ubx/
abdA
PG7
hox7/antp
PG6
hox6/lox5/
ftz
PG5
hox5/scr
PG4
hox4/dfd
PG1
hox1/lab
PG3
hox3/zen
PG2
hox2/pb

**Figure S3.** Orthology analysis of *Spadella cephaloptera* Hox genes. The phylogenetic tree of Hox genes is based on bilaterian protein sequences obtained from published literature and BLAST searches of the NCBI GenBank. The tree was generated using Maximum Likelihood analysis and SH-aLRT test implemented in IQTREE with the following configurations: VT+F+I+G was selected based on ModelFinder and SH-aLRT and ultrafast bootstrap set to 1000. The support values of branches indicate SH-aLRT and ultrafast boostrap support (SS% / UFBS%). The tree is rooted with *even-skipped* (*evx*) sequences from different bilaterian representatives. Each paralogous group (PG) is highlighted in different box colors and *S. cephaloptera* Hox gene sequences are in yellow boxes. Species abbreviations are in Table S1.

```
                  10        20        30        40        50        60
          ----:----|----:----|----:----|----:----|----:----|----:----|----:--
Consensus PRRLRTAYTNTQLLELEKEFHFNKYLCRPRRIEIAASLDLTERQVKVWFQNRRMKHKRQTQKKKXXD
Acr_Hox2  T......................................................Y...S.IQ.HG-
Sva_Hox2  S......................................................Y...S.SGRSKS
Mal_Pb    ..........S............................................Y...SG------
Cte_Pb    .......................................................F....G.GSGNS
Csa_Pb    ............................................................SVM.DD.
Ctu_Pb    -------------------............................--------------------
Tca_Mxp   ............................................................LG.QGD.
Eka_Pb    ...........................................................NLG.GAE.
Pca_Pb    --------------------....................S....----------------------
Sce_Hox2  ..........................................................HSLQTGPGA
Bpl_Hox2  H..I......N.........FN.........V...SN.S.......I..........KERSH..NRK
Bla_Hox2  S.....VF.............Y...V.K...K...SF...N.....I.......RQ..RDT.SRSEI
Sko_Hox2  H..V...F.............Y.............SM...S..................IM.AAVSG
Hsa_HoxA2 S..............................V....L........................C.ENQN
Hsa_HoxB2 A..............................V....L........................HREPP.
```

**Figure S4.** Hox domain alignment of bilaterian PG2**.** Diagnostic residues for the Hox2/Pb are highlighted in yellow. Chaetognath sequences are in dark blue and *Spadella cephaloptera* sequences are in **bold** letters. Species abbreviations are in Table S1.

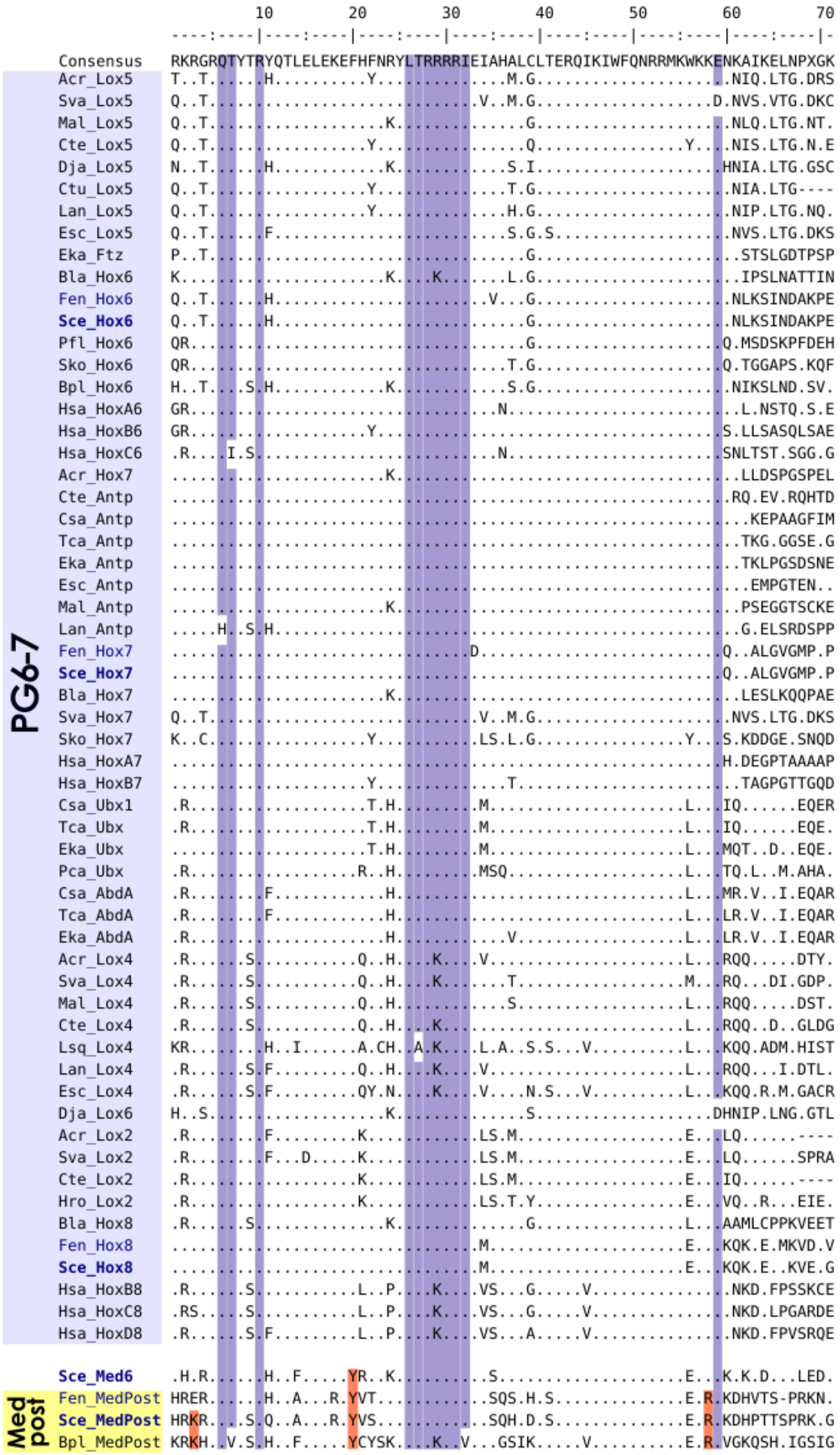


**Figure S5.** Hox domain alignment of bilaterian PG6-7 genes (light blue) and medpost genes (yellow) from rotifer and chaetognaths**.** Diagnostic residues for the central class and posterior Hox are highlighted in purple and orange, respectively. Chaetognath sequences are in dark blue and *Spadella cephaloptera* sequences are in **bold** letters. Species abbreviations are in Table S1.

```
                          10        20        30        40        50        60        70
              --------:----|----:----|----:----|----:----|----:----|----:----|----:----|-
Consensus       RSGRKKRKPYTKYQTLELEKEFLFNMYITRZRRWELSRXLNLTERQVKIWFQNRRMKXKKLNEREKAEQKENP
Csa_AbdB        VTV.......S.F............A.VSKQK....A.N..................S..TSQ.NAENNQN.T
Eka_AbdB        VTV.......S.F............A.VSKQK....A.N..................N..N.Q.NLENN.---
Tca_AbdB        VTV.......S.F............A.VSKQK....A.N..................N..NSQ.QA.Q.QN.N
Pca_AbdB        V.V......................A.VSKQK....A.T..................S..S.QK.TEK.RQHQ
Pca_HB4         SFV.S.....S.Q.IVD....YKV.P....Q..F.MAHN.G.S..............T..HLLTLRV------
Sro_Post        NVS....R..P.N............T....E..L.IA.S....D.............N..QHSGGPGMGPPI.
Acr_Post2       PK.........R...MV..N...N.S....QK...I.CK.Q.......V........R......A..QI..ES
Cte_Post2       PKQ........R...MV..N..IN.S....QK...I.CK.H.S.....V........R......A.SLI.DHE
Esc_Post2       TK.........R...MV..N...NSS....QK...I.CK.Q.......V........R......A..RLR.DR
Lan_Post2       ------.....R...MV..N...N.A....QK...I.CK.H.S.....V........R......A..LF.SED
Mal_Post2       PRT........R...MV..N..MT.S....QK...I.CK.H.......V........R....S.A.VKSEVEN
Ctu_Post2       S.S........R...MV..T..IN.S....QK...I.CR.R.......V........R....D.A.NA.LTVQ
Cte_Post1       VNPK......S.P.VSA..N.YSTST...KA..K.VA.E.D.....I...Y....I.E..IATKRAKV.SDLF
Esc_Post1       IAL..R.R..S...IA...R.YALST..SKS......QL...S...I........I.A...QK.DETLKTQT.
Sva_Post1       VTL..R.R..S.F.IA...R.Y-NGS.VSES......QLI..S...I........I.A..IIK.DDISPQVSH
Lan_Post1       IHM.......S...IA...R.YVS.T..SKPK.....QR.Q.S..............E..VKGGKQT------
Fen_PostA       ---.........H..FI..Q.Y.MST....Q..L..A.N.S................T...R..N.VSVGPDS
Sce_PostA       TI..........H..FI..Q.Y.MST....Q..L..A.N.S................T...R..N.VSVGADS
Fen_PostB       GKC.T.....E.WV.YL..E.Y.S.T...KQK.Y...YRTS................S...R..STSGATG.E
Sce_PostB       GKC.T.....E.WV.YL..E.Y.S.T...KQK.Y...YRTS................S...R..STSGGPDVD
Sce_PostC       S......V...RS..GW..R..VM.T....K..V..AYM..................Q...M...RSKHHHHH
Sce_PostD       PGQ....V..SRF.IK...R.Y.R.T....K..L..AYF.S................S.R.L..Q.KAAEWGA
Pfl_Hox9-10     A......C....F..............L..E..V.I..L..................L..Q.H.NTICMIH--
Sko_Hox9-10     A......C....F..............L..E..VDIA.L..................L..Q.Q.NATMLH---
Bla_Hox9        H.S....C...RF..........Y...L..E..Y.I.QHV..S..............M..MSKQRQEQ.QP..
Hsa_HoxA9       ..T....C....H..............L..D..Y.VA.L..................M..I.KDRAKD-----
Hsa_HoxB9       ..S....C...................L..D..H.VA.L...S..............M..M.KEQGK.-----
Hsa_HoxC9       ..T....C...................L..D..Y.VA.V..................M..M.KEKTDKEQS--
Hsa_HoxD9       ..T....C...................L..D..Y.VA.I..................M..MSKEKCPK-----
Bla_Hox10       .V.....C......I............VS.E..Q.I..HV..SD.............M.RM.KAREEQIRNHQ
Hsa_HoxC10      K......C....H..............L..E..L.I.KTI...D.............L..M.RENRIRELTSN
Hsa_HoxD10      K......C....H..............L..E..L.I.KSV...D.............L..MSRENRIRELTAN
Bfl_Hox11       K.T....C..................FV..E..Q.IA.Q....D.............M.RMKQ.AMQQLM.EK
Hsa_HoxA11      QRT....C......IR...R..F.SV..NKEK.LQ...M....D.............E..I.RDRLQYYSA..
Hsa_HoxD11      QRS....C......IR...R..F..V..NKEK.LQ...M....D.............E....RDRLQYFTG..
Hsa_HoxC11      PRT....C..S.F.IR...R..F..V..NKEK.LQ...M....D.............E...SRDRLQYFSG..
Pfl_Hox11-13a   TRN....................Y...L..D..TDI..A..................L..MRL..EN.R.QQE
Pfl_Hox11-13b   TPR.T..R..S.M.IY....A.QQ.A.L..E..QKY.QQ..................S..QV....M.E..RQ
Pfl_Hox11-13c   TPR.T..R..S.L.IF......QQ...L..D..TR.AQT..................L..MT...HS..EVLH
Sko_Hox11-13a   TRN....................Y...L..D..TDIA.A...S...I..........L..MRL..EN.R.QQD
Sko_Hox11-13b   TPR.T.......M.IF...QA.QN...L..E..TK..QQ.S.S...I..........L..MT....L.E..LR
Sko_Hox11-13c   TPR.T..R..S.L.IF......QQ...L..D..SR..QA.......I..........L..MTD..RN..DMMQ
Bla_Hox12       Q.S....C..S.V.L........Y......EQ.G.IA.KV...D.............M.RMKQ.HEE.AFRAH
Hsa_HoxC12      SRS.......S.L.LA...G...V.EF...Q..R...DR...SDQ............K.R.LL..Q.LSFF--
Hsa_HoxD12      GEA.........Q.IA...N...V.EF.N.QK.K...NR...SDQ............K.RVVL..Q.LALY--
Bla_Hox13       .G.....C..S...LSV..Q.YIQ.R.VS.ET.L...QR....D.............Q.R.EF.SGNQT----
Hsa_HoxA13      .R.....V....V.LK...R.YAT.KF..KDK.RRI.ATT..S....T.......V.E..VINKL.TTS----
Hsa_HoxB13      .R.....I..S.G.LR...R.YAA.KF..KDK.RKI.AATS.S...IT.......V.E..VLAKV.NSATP--
Hsa_HoxC13      .R.....V....V.LK.....YAASKF..KEK.RRI.ATT..S....T.......V.E..VVSKS..PHLHST
Hsa_HoxD13      .R.....V....L.LK...N.YAI.KF.NKDK.RRI.AAT..S....T.......V.D..IVSKL.DTVS---
Bla_Hox14       KPV.P..R..S...LN...N.YVQ.Q..S.DK.LQ..QK................I.Q...DR.NS-------
Bla_Hox15       PRT....R..S.P.LAL..D.YASQKFL.KEK.K.I.ESSS.S....M.........E...AR.AA.QRHAQH
```

**Figure S6.** Hox domain alignment of bilaterian PG9 – 14 genes**.** Diagnostic residues for the posterior class Hox are highlighted in orange. Chaetognath sequences are in dark blue and *Spadella cephaloptera* sequences are in **bold** letters. Species abbreviations are in Table S1.

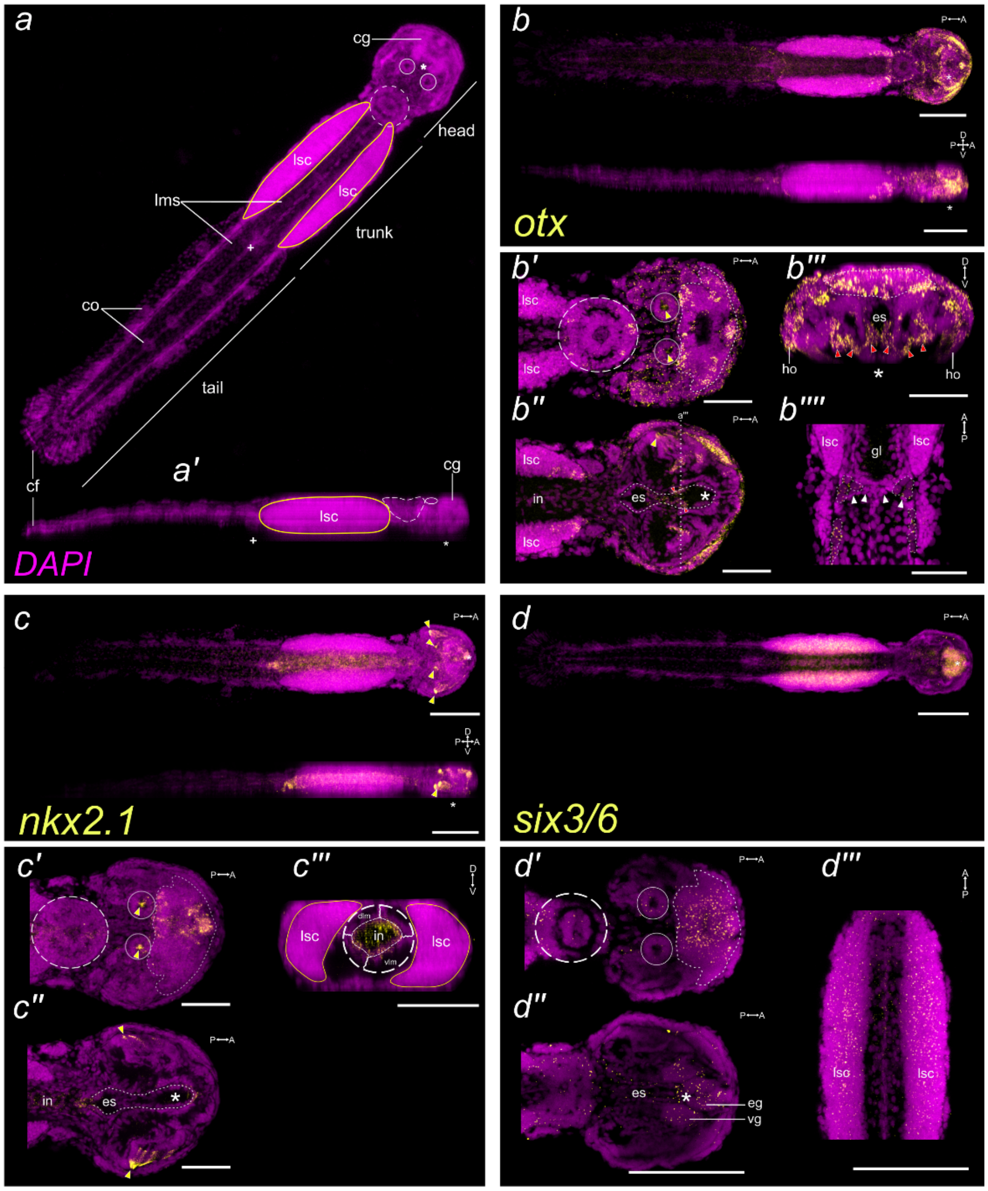


**Figure S7.** Expression pattern of *Sce-otx, Sce-nk2.1,* and *Sce-six3/6* in early juveniles (7-10 dph) of *S. cephaloptera*. Gene transcripts (in yellow) are visualized with AlexaFluor647 (b, c) or AP-Fast Blue (d) and nuclei with DAPI (purple). General morphology of a *S. cephaloptera* early juvenile in maximum intensity projection of the whole animal in DAPI. Eye locations are encircled and corona ciliata in a dashed circle. The asterisk and crosshair indicate the position of the mouth and anal opening, respectively. Top (a) and bottom (a') figures are coronal and lateral view, respectively. (a') Lateral view of the early juvenile. (b – b'''') *Sce-otx* expression pattern. (b') Coronal profile of the dorsal-most structures of the head. (b'') Coronal profile exposing the mouth opening and esophagus (es). (b''') Transverse profile along the posterior portion of the cerebral ganglion (cg). (b'''') Higher magnification of the dividing germ cells (dotted outline). Female (top) and male germ cells (bottom) are separated by the trunk-tail septum (arrowheads). (c – c''') *Sce-nk2.1* expression pattern. (c') Coronal profile of the anterior most region of the head. (c'') Coronal section showing expression at the posterior esophagus (es) and anterior mouth. (c''') Transverse section of the trunk showing broad expression in the intestine (in). (d – d''') *Sce-six3/6* expression pattern. (d') Coronal profile of the anterior region. (d''). Coronal profile exposing the vestibular (vg) and esophageal ganglia (*eg*). (d''') Coronal profile of the trunk showing dispersed expression of *Sce-six3/6* in the lateral somata clusters (*lsc*). Scale bars: 50 µm, except b, c, and d (100 µm). Yellow arrowheads indicate autofluorescence, which are typically observed in the pigmented region of the eyes and the grasping spines. cf, caudal fin; cg, cerebral ganglion; co, ciliary tuft/fence organ; es, esophagus; in, intestine; lms: longitudinal mucles somata; lsc, lateral somata clusters.

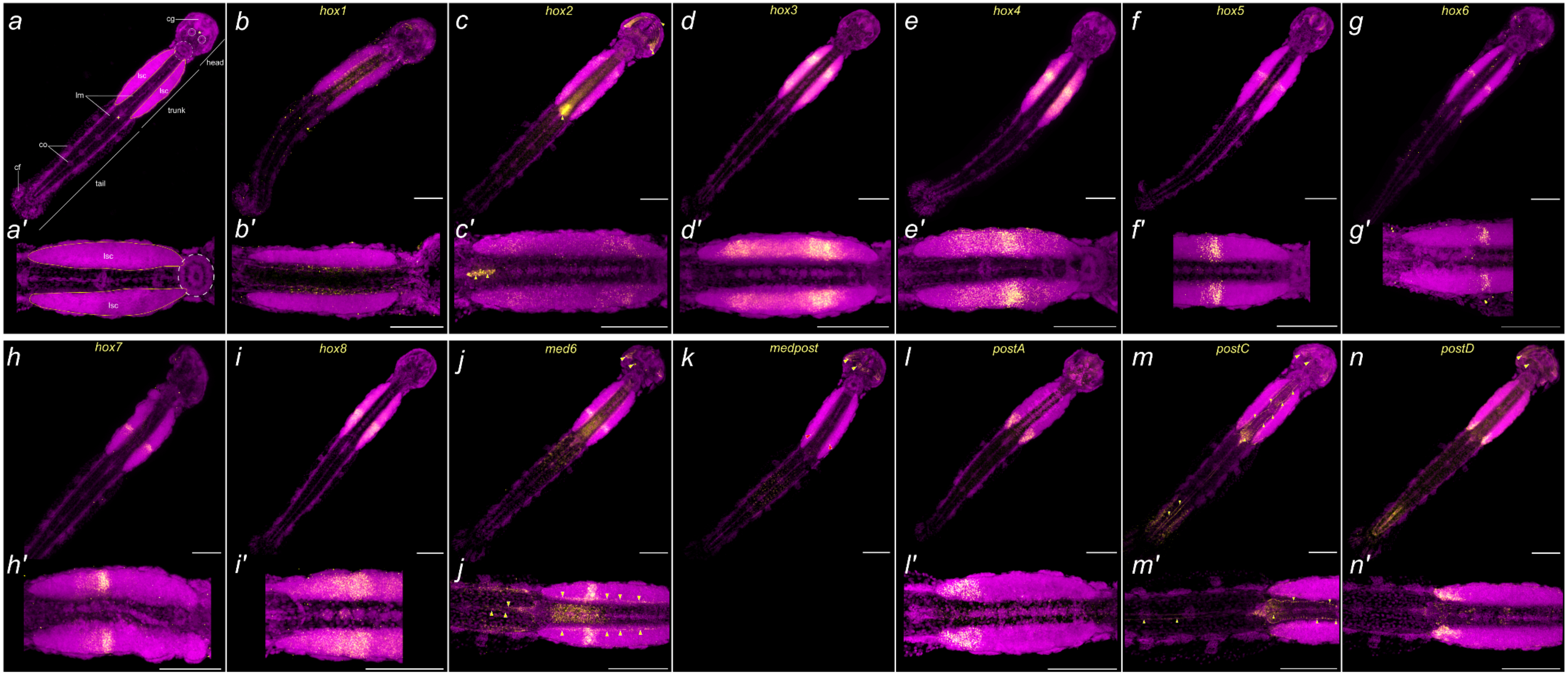


**Figure S8.** Expression pattern of *Hox* genes in early juveniles (7-10 dph) of *S. cephaloptera*. Gene transcripts (in yellow) are visualized with AP-Fast Blue (b – i, l) or AlexaFluor647 (c, j, k, m, n) and cell nuclei with DAPI (purple). (a) An early juvenile showing the sections of head, trunk, and tail. Eye locations are encircled and corona ciliata is in a dashed circle. The asterisk and crosshair indicate the position of the mouth and anal opening, respectively. (a') A magnified section of the trunk. (b - n) Coronal maximum projection of the whole animal. (k) The expression of *Sce-medpost* in the lateral somata clusters (*lsc*) is very weak (red arrowheads). Yellow arrowheads indicate autofluorescence, which are typically observed in the pigmented region of the eyes and the grasping spines, or non-specific staining. Scale bars: 100 µm. cf, caudal fin; cg, cerebral ganglion; co, ciliary tuft/fence organ; longitudinal muscles ; lm, lateral somata clusters.
